# Uncovering the essential roles of human GCP 2 orthologs in *Caenorhabditis elegans*

**DOI:** 10.1101/2023.02.27.529682

**Authors:** Lucie Panska, Stepanka Nedvedova, Vojtech Vacek, Daniela Krivska, Lukas Konecny, Filip Knop, Zsofia Kutil, Lubica Skultetyova, Adrian Leontovyc, Lenka Ulrychova, Judy Sakanari, Masako Asahina, Cyril Barinka, Marie Macurkova, Jan Dvorak

## Abstract

Human glutamate carboxypeptidase 2 (GCP2) from the M28B metalloprotease group is an important target for therapy in neurological disorders and an established tumor marker. However, its physiological functions remain unclear. To better understand general roles, we used the model organism *Caenorhabditis elegans* genetically manipulate its three existing orthologous genes and evaluate the impact on worm physiology. The results of gene knockout studies showed that *C. elegans* GCP2 orthologs affect the pharyngeal physiology, reproduction, and structural integrity of the organism. Promoter-driven GFP expression revealed distinct localization for each of the three gene paralogs, with *gcp-2*.*1* being most abundant in muscles, intestine, and pharyngeal interneurons, *gcp-2*.*2* restricted to the phasmid neurons, and *gcp-2*.*3* located in the excretory cell. This study provides new insight into the unique phenotypic effects of GCP2 gene knockouts in *C. elegans*, and the specific tissue localizations. We believe that elucidation of particular roles in a non-mammalian organism can help to explain important questions linked to human GCP2 physiology and in extension to GCP2 involvement in pathophysiological processes.

## Introduction

The M28 protease family is an intensively studied group of proteolytic enzymes mainly due to its significance in human pathologies (MEROPS) (Rawlings *et al*, 2018). Most of the research attention is devoted to the M28B subfamily because of its importance in pathological processes in humans. However, members of this subfamily are ubiquitously expressed across all phyla, from unicellular organisms to plants and animals. The unique feature of this di-zinc metalloprotease group is the presence of enzymes with either amino- or carboxypeptidase activity (Rawlings *et al*, 2018). In the human genome, five genes encoding M28B paralogs possessing carboxy- or amino-peptidase activities were identified to date with either partially known or completely unknown functions (Rawlings *et al*, 2018; Tykvart *et al*, 2015b; Hlouchova *et al*, 2012). Interestingly, transferrin receptors with non-enzymatic function (transferrin receptor protein and transferrin receptor 2 protein) are also structurally related to the common ancestor together with M28B family proteases (Lambert & Mitchell, 2007). The most studied human protease from the M28B subfamily is glutamate carboxypeptidase 2 (GCP2 or GCPII; EC 3.4.17.21), which is also known as a prostate-specific membrane antigen (PSMA), N-acetylated-alpha-linked acidic dipeptidase (NAALADase), and folate hydrolase 1 (FOLH1) (Lambert & Mitchell, 2007; Tykvart *et al*, 2015a).

In humans, GCP2 is a membrane-bound metallopeptidase composed of zinc-dependent homodimers (Pavlicek *et al*, 2012). This transmembrane metalloprotease comprises three structural domains: protease, apical, and C-terminal, which all contribute to the architecture of the bimetallic active site (Pavlicek *et al*, 2012). In addition to zinc ions, human GCP2 also requires Ca^2+^ ions for proper folding, stability, and enzymatic activity (Ptacek *et al*, 2018). GCP2 is primarily expressed in the nervous system (astrocytes and Schwann cells), kidney (Matteucci *et al*, 2017), and jejunal brush membranes. Lower expression levels were also reported in salivary and lacrimal glands, prostate, heart, pancreas, bladder, skin, breast, liver, lung, colon, and testis (Rovenská *et al*, 2008; Silver *et al*, 1997). Despite well-described expression profiles in human organs, there is an absence of data on GCP2 physiology in most tissues, except for the nervous system and small intestine. In the nervous system, GCP2 is involved in communication between neurons and glial cells by hydrolyzing N-acetyl-aspartyl-glutamate (NAAG), the most abundant neuropeptide in the mammalian brain. In the digestive system, GCP2 plays an important role in the absorption of dietary folates as it cleaves off their C-terminal poly-glutamylated tails, thus enabling the absorption of folate mono-glutamate into the bloodstream (Visentin *et al*, 2014). In addition to their function as hydrolases/peptidases, several reports allude to a non-proteolytic role(s) of GCP2, such as the roles associated with the anaphase-promoting complex in prostate cancer cells or to activation of the NF-κB signaling pathway in cell proliferation (Rajasekaran *et al*, 2005). Human GCP2 is an attractive target for pharmacological interventions in neurodegenerative diseases and cancer. Understanding and dissecting potentially diverse biological functions of the enzyme is therefore desirable. Consistent with the role of GCP2 (NAAG hydrolysis), changes in the NAAG levels and/or changes of GCP2 enzymatic activities correlate with pathologic conditions, including Alzheimer’s disease, Huntington’s disease, amyotrophic lateral sclerosis (Tsai *et al*, 1991; Plaitakis, 1990), epilepsy, schizophrenia (Fricker *et al*, 2009), and stroke (Šácha *et al*, 2007). The inhibition of GCP2 activity has already been proven as an effective neuroprotective therapy against ischemic brain injury in animal models (Lu *et al*, 2000). GCP2 expression has also been found in the tumors of the neovasculature (Evans & Blumenthal, 2000), kidney (Maurer *et al*, 2016), digestive tract (Haffner *et al*, 2009) and in various subtypes of bladder cancer (Samplaski *et al*, 2011). Other members of the M28B family in human are PSMA-L, glutamate carboxypeptidase 3 (GCP3), NAALADase L, and NAALADase L2. Their physiological roles are either partially described or not clarified yet (Pavlicek *et al*, 2012; Bařinka *et al*, 2012). However, NAALADase L was recently characterized as an enzyme with aminopeptidase activity expressed in intestinal tissue (Tykvart *et al*, 2015a), and GCP3 was shown to have overlapping substrate specificity and inhibition profile with GCP2 (Bacich *et al*, 2002; Hlouchova *et al*, 2009).

The studies of the physiological functions of this protease group are complicated for many reasons. The most notable obstacle is the presence of several paralogs in a single organism as seen in mammals including humans, therefore more phylogenetically basal model organisms may offer significant advantages. During evolution, a wide spectrum of M28B proteases evolved by duplications and subsequent speciation occurred as evident from the example of the aforementioned human paralogs (Lambert & Mitchell, 2007). In contrast to higher organisms, lower organisms (including nematodes) possess only a single or few genes coding M28B metalloproteases (Howe *et al*, 2017), thus offering opportunities to study ancestral proteases before their diversification. So far, none of the GCP2 orthologs from lower basal organisms has been intensively studied. For that reason, we selected the free-living nematode *Caenorhabditis elegans* as a model to investigate physiological functions of GCP2 orthologs. *C. elegans* serves as an excellent laboratory model in a variety of research areas due to a well-characterized genome, relatively simple body organization, uncomplicated maintenance in laboratory conditions, and the existence of a host of experimental tools to study its physiology, including transgenic approaches (Riddle *et al*, 1997). We focused on all three genes coding subfamily M28B orthologs named CeGCP2.1, CeGCP2.2, and CeGCP2.3 encoded by *gcp-2*.*1, gcp-2*.*2*, and *gcp-2*.*3* genes, respectively (Harris *et al*, 2020). Their nomenclature however is without any known relevance to the substrate preferences or the enzymatic mode of actions and it is solely based on existing gene annotations (Davis *et al*, 2022).

In our study, we report the most pronounced phenotypes observed in a member of this metalloprotease group in any animal model to date. After performing a wide range of phenotypic studies, we identified several unique phenotypes that, together with their localization and expression, led to the unexpected findings that all three genes acquired distinct and unique functions.

## Results

### Primary sequence alignments and homology modeling of *C. elegans* M28 metalloproteases

Based on a database search (WormBase, genome assembly WS284), all three genes are located on the X chromosome. *Gcp-2*.*1* gene is located in the X: 304131..305653 chromosomal region. And the *gcp-2*.*2* and *gcp-2*.*3*, are localized adjacent to each other in the X:11537714..11541538 and X:11542138..11545061 regions, respectively (Davis *et al*, 2022). The corresponding proteins are termed CeGCP2.1, 2.2, and 2.3, and this respective terminology will be used throughout this report. All orthologs share the identical domain organization shown in Fig. 1, which corresponds to their human GCP2 ortholog (hsGCP2) (also later verified by homology modeling, Fig. 2) consisting of transmembrane, protease, apical, and C-terminal dimerization domains (Barinka *et al*, 2008). Interestingly, a deletion of 45 amino acids was observed for CeGCP2.3 in C-terminal dimerization domains that could negatively influence the quaternary homodimeric structure of this isoform (Fig. 1). Furthermore, three splice variants exist for CeGCP2.1, where the CeGCP2.1b (751 amino acids) isoform lacks the sequence encoding the intracellular part (as compared to the CeGCP2.1a isoform; 770 amino acids), and the shortest CeGCP2.1c isoform (576 amino acids) additionally lacks the C-terminal dimerization domain (Fig. EV1). The results of the protein structure analysis showing sequence identity and similarity of hsGCP2 and CeGCP2 orthologs are presented in Table 1.

**Table 1:** Overview of the protein sequence identity and similarity of hsGCP2 and CeGCP2 orthologs.

| Identity [%] |  |  |  |  | Similarity [%] |  |  |  |
| --- | --- | --- | --- | --- | --- | --- | --- | --- |
|  | hsGCP2 | GCP2.1 | GCP2.2 | GCP2.3 | hsGCP2 | GCP2.1 | GCP2.2 | GCP2.3 |
| **hsGCP2** | 100.00 | 29.73 | 27.58 | 24.90 | 100.00 | 48.28 | 43.83 | 41.90 |
| **GCP2.1** | 29.73 | 100.00 | 34.13 | 28.43 | 48.28 | 100.00 | 52.64 | 44.54 |
| **GCP2.2** | 27.58 | 34.13 | 100.00 | 57.25 | 43.83 | 52.64 | 100.00 | 66.20 |
| **GCP2.3** | 24.90 | 28.43 | 57.25 | 100.00 | 41.90 | 44.54 | 66.20 | 100.00 |

**Figure 1.**
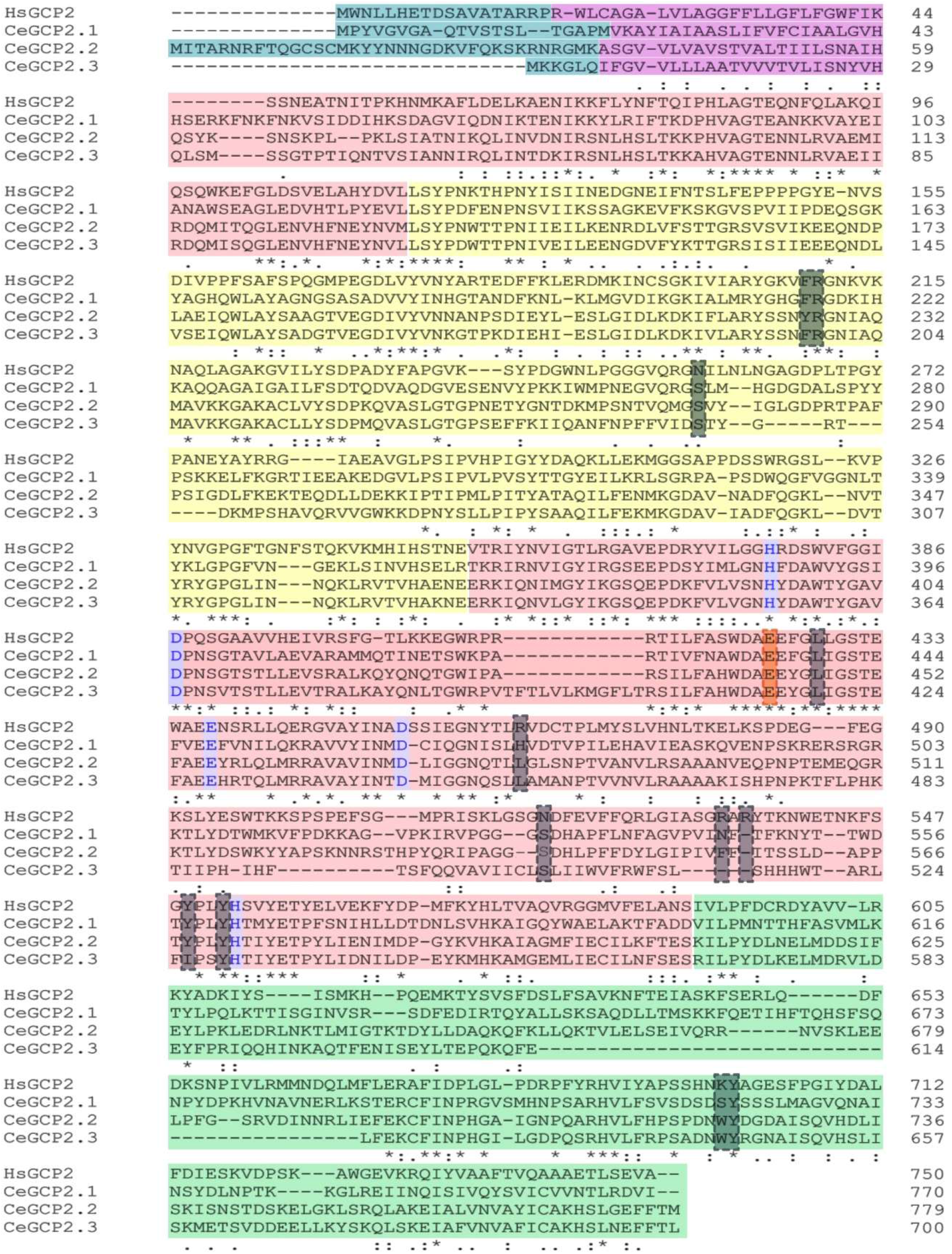
Primary sequence alignment of CeGCP2.1, CeGCP2.2, CeGCP2.3, and hsGCP2. The intracellular and transmembrane parts of GCP2 are shaded turquoise and magenta, respectively. Protease-like, apical, and C-terminal dimerization domains are shaded red, yellow, and green, respectively. The catalytic acid/base glutamate is marked by red framing with red shading and residues coordinating the active-site zinc ions are marked with a purple frame. Residues delineating specificity pockets are indicated by a gray box with gray shading The alignment and subsequent identity and similarity analyses are shown in Table. 1.

**Figure 2.**
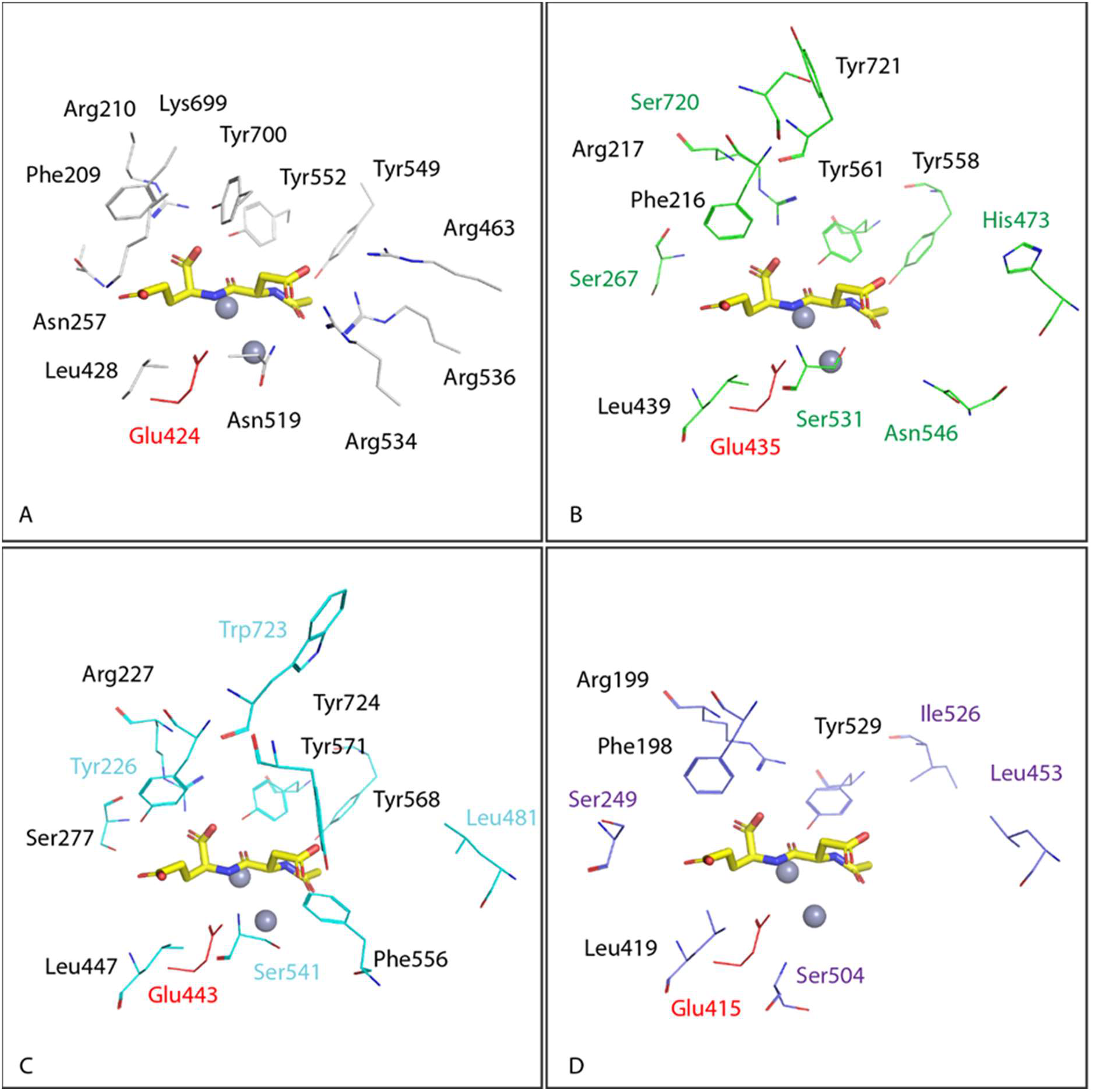
Comparison of active sites of hsGCP2 and CeGCP2 orthologs. A Residues delineating the substrate-binding pocket of hsGCP2, are shown in line representation, with atoms colored red, blue, and gray for oxygen, nitrogen, and carbon, respectively. Active-site bound NAAG is shown in stick representation with carbon atoms colored yellow. Zinc ions are shown as gray spheres. Residues in CEGCPs that are structurally equivalent to hsGCP2 ortholog are shown in line representation. B, C, D Conserved residues have black labels (B) (C) (D), while non-conservative substitutions are colored green, cyan, and violet for CeGCP2.1 (B), CeGCP2.2 (C), and CeGCP2.3 (D), respectively. The critical proton shuttle glutamate residue is highlighted in red. The figure was generated using PyMol 2.4.1.

To provide structural insight into CeGCP2 variants, we prepared homology models of all three *C. elegans* enzymes and compared them with the X-ray crystal structure of the hsGCP2 (PDB code: 3BXM (Klusák *et al*, 2009); Fig. 2). In line with the sequence alignment, our homology models reveal the conservation of residues coordinating Zn^2+^ ions in the bimetallic active site of all proteins. Furthermore, Glu424, the proton shuttle residue critical for the catalytic activity of hsGCP2 (Mesters *et al*, 2006), is also conserved in all orthologs. As both zinc ions and the proton shuttle are essential for GCP2 folding and hydrolytic activity, it is fair to speculate that *C. elegans* orthologs can be enzymatically competent.

Contrary to absolute structural conservation of the zinc-dependent active site, marked differences are observed for residues delineating substrate specificity pockets in the internal cavity of these enzymes, pointing towards their distinct substrate specificities. For example, hsGCP2 has a preference for substrates that are negatively charged at both P1 and P1’ positions (Barinka *et al*, 2002). At the P1’ position, specificity for glutamate residues is achieved by the intricate network of hydrogen bonds and most notably the presence of positively charged Lys699 (Mesters *et al*, 2006; Navrátil *et al*, 2016; Pavlicek *et al*, 2012; Ferraris *et al*, 2012). In *C. elegans* orthologs, the Lys699 is substituted by Ser720, Trp723, and Trp644 in CeGCP2.1, 2.2, and 2.3, respectively. Similarly, the arginine patch comprised of side chains of Arg463, Arg534, and Arg536 is critical for the recognition of negatively charged residues in the non-prime specificity pocket of hsGCP2 (Barinka *et al*, 2008), yet these residues are missing in studied orthologs and/or are replaced by polar/hydrophobic amino acids (Figs. 1 and 2). Taken together, GCP2 orthologs are likely proficient hydrolases with substrate specificities different from the human enzyme, and further studies are warranted to identify their physiological substrates in *C. elegans*.

### Phylogenic diversification of M28B metallopeptidases within nematodes

To analyze phylogenetic relationship of M28B peptidases in nematodes, we created a maximum likelihood phylogenetic tree (Fig. EV2). Here, CeGCP2.1 sits in a well-supported clade with other orthologous sequences of the genus *Caenorhabditis*. However, CeGCP2.2 and 2.3 homologs form a clade that branches at the base of group A. This position in the phylogenetic tree shows that CeGCP2.2 and 2.3 are phylogenetically relatively distant from GCP2.1. Nonetheless, they are closely related to each other and most likely originated from gene duplication. These findings suggest that CeGCP2.2 and 2.3 arose by later diversification from the CeGCP2.1. As seen from the gene databases (Davis *et al*, 2022), the *gcp-2*.*1* gene encodes an M28B metalloprotease ancestral for nematodes, while any other gene duplications or modifications result from an adaptive response to the environmental conditions.

### Relationship in the gene expression levels of the three *C. elegans* M28B peptidases

To get insight into the physiological roles of M28B peptidases, we first analyzed their gene expression by means of RT-qPCR and then explored the effect of knocking out the *gcp-2*.*1* (R57.1; allele name – ok1004), *gcp-2*.*2* (C35C5.2; allele name – tm6541), or *gcp-2*.*3* (C35C5.11; allele name – tm5414) genes (Appendix Fig. S1) on each other’s expression (Fig. 3A and 3B). All three genes including *gcp-2*.*3*, which is annotated as a pseudogene in WormBase, are expressed (Fig. 3A). *Gcp-2*.*1* is the most abundant transcript and *gcp-2*.*3* has the lowest level of expression in wild-type worms. Based on RT-qPCR data, the expression of the *gcp-2*.*2* gene was not affected in *gcp-2*.*1* mutant animals, while the expression of *gcp-2*.*3* was increased approximately 2-fold in comparison to the wild-type population. The knockout of *gcp-2*.*2* completely suppressed the expression of the *gcp-2*.*3* gene to only 1.8% of the wild-type levels. *Gcp-2*.*3* knockout has significantly reduced the expression of *gcp-2*.*1* and *gcp-2*.*2* genes 8- and 4-fold, respectively, when compared to the wild-type control (Fig. 3B). For those reasons, the impacts of the particular knockouts on the expression must be considered while investigating specific effects on the worm phenotypes, as shown below.

**Figure 3.**
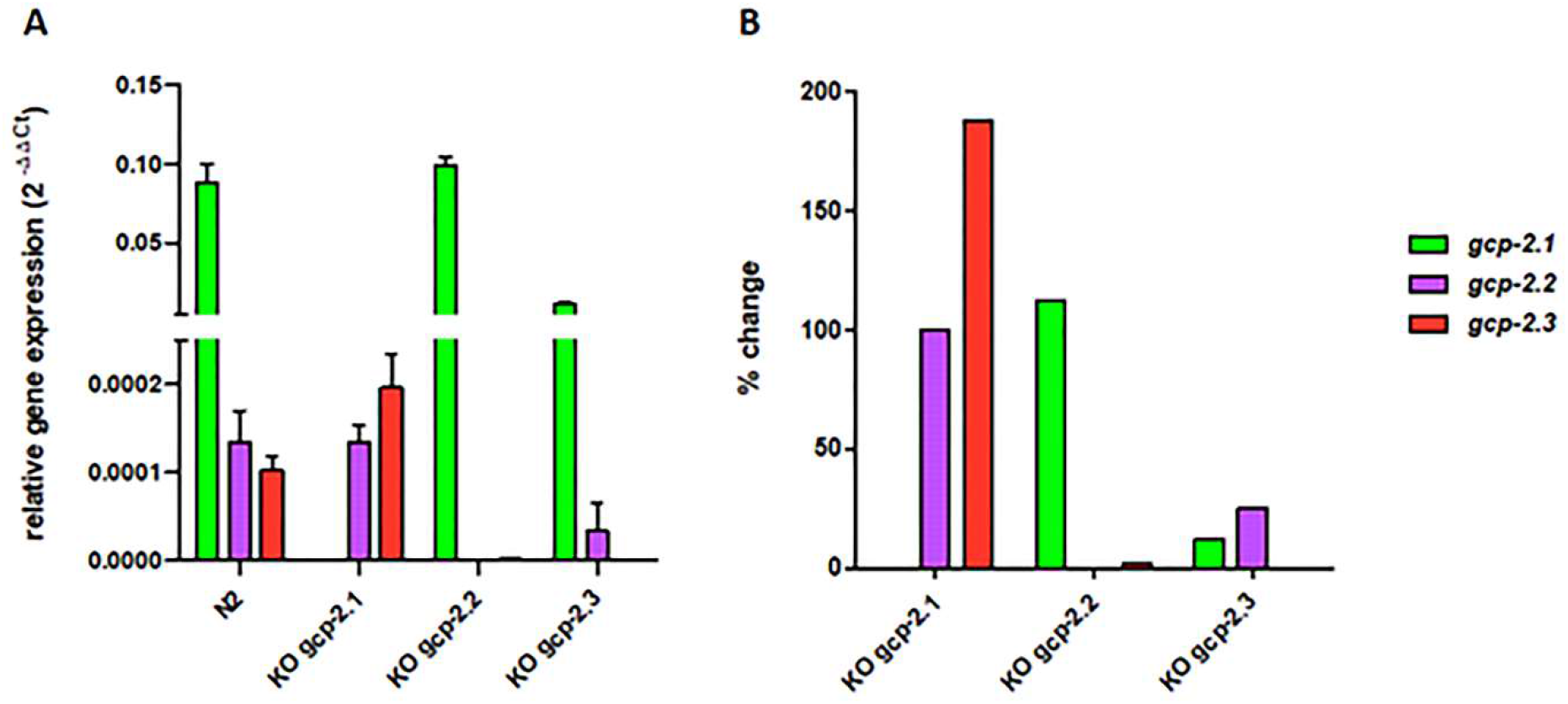
Gene expression of M28B metalloproteases in adult *C. elegans* evaluated by RT-qPCR. A *gcp-2*.*1, gcp-2*.*2*, and *gcp-2*.*3* gene expression in wild-type N2 strain and mutants B Relative gene expression in the particular mutants compared to the expression in wild-type worms. Quantification of *gcp-2*.*1* (green), *gcp-2*.*2* (purple), and *gcp-2*.*3* (red) gene expression levels were performed using RT-qPCR; cycle threshold (Ct) values normalized to *tba-1* as a housekeeping gene, are shown as mean +/-standard deviation.

### Tissue-specific expression of CeGCP2

To determine expression sites of the three M28B paralogs *in vivo*, we used standard promoter-driven GFP (green fluorescent protein) expression. Upstream regions of each gene used to generate transgenes are described in Fig. S11. Our data revealed that expression patterns of the three proteases differ markedly, and their expression is limited to distinct tissue structures. Consistent with the high expression of the *gcp-2*.*1* gene (Fig. 3 A), high level of extrachromosomal transgene *gcp-2*.*1p::GFP* expression was detected in several tissues of the wild type worms, including the intestine, somatic muscles (Fig. 4 D), and the nervous system of adult worms (Fig. 4 E, F, G, H). Specifically for the nervous tissue, the *gcp-2*.*1p::GFP* is mainly expressed in class SAA pharyngeal interneurons (a paired motor neuron localized in the head that uses acetylcholine signaling) and their anterior processes (Fig. 4 F, G, H). L3 larvae, in contrast, displayed expression of *gcp-2*.*1p::GFP* predominantly in the muscles (Fig. 4 B). Lower levels of *gcp-2*.*2* and *gcp-2*.*3* gene expression (Fig. 3 A) correspond to the restricted GFP expression pattern, where each paralog is observed only in a single site: phasmid neurons (PHAL/R, PHBL/R) and their processes (Figs. 5 B, C) in adults and L3 larvae (Fig. EV3 B) for *gcp-2*.*2p::GFP* and the excretory cell and its extended bilateral canals reaching from the nose to the tail region of the adult (Fig. 6 B) and larvae (Fig. EV3 D) for *gcp-2*.*3p::GFP* (for an overview see Fig. 9).

**Figure 4.**
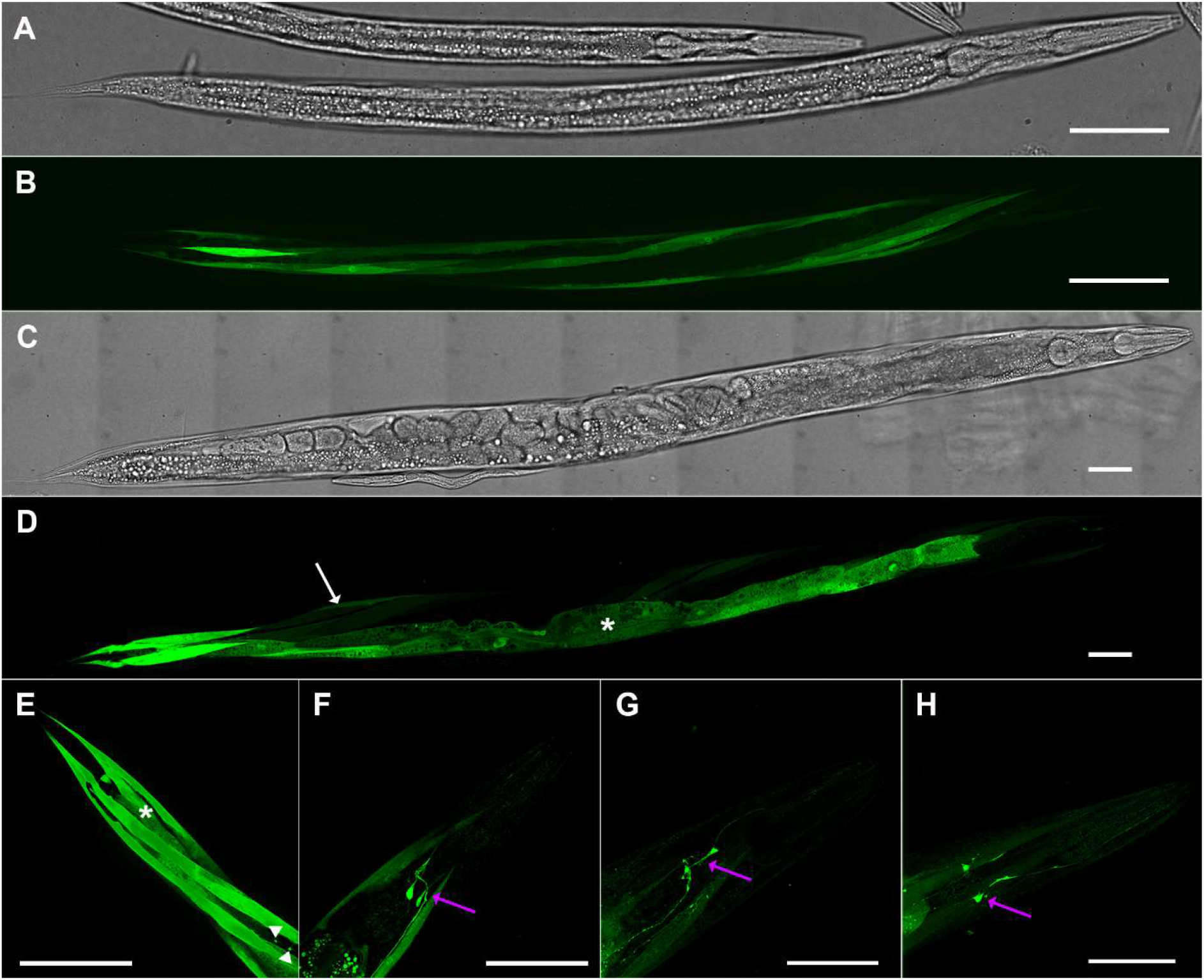
Expression pattern of *gcp-2*.*1* in the wild-type *C. elegans* visualized by GFP. A Bright-field image of L3 larvae. B *gcp2*.*1p::GFP* expression was localized to somatic muscles of L3 larvae. C Bright-field of adult worm. D Strong expression of *gcp-2*.*1p::GFP* was observed as well in the adult intestine (asterisk), and somatic muscles (arrow). E Detailed localization of the *gcp-2*.*1* in the posterior part of the body of wild-type adult *C. elegans*. The expression is visible in the intestine (asterisk), muscles, and slender muscle arms reaching the nervous cord (arrowheads). F, G, H Expression of *gcp-2*.*1p::GFP* in the class SAA pharyngeal interneurons and their anterior processes (purple arrows) of adult *C. elegans*. Scale bars represent 50 µm.

**Figure 5.**
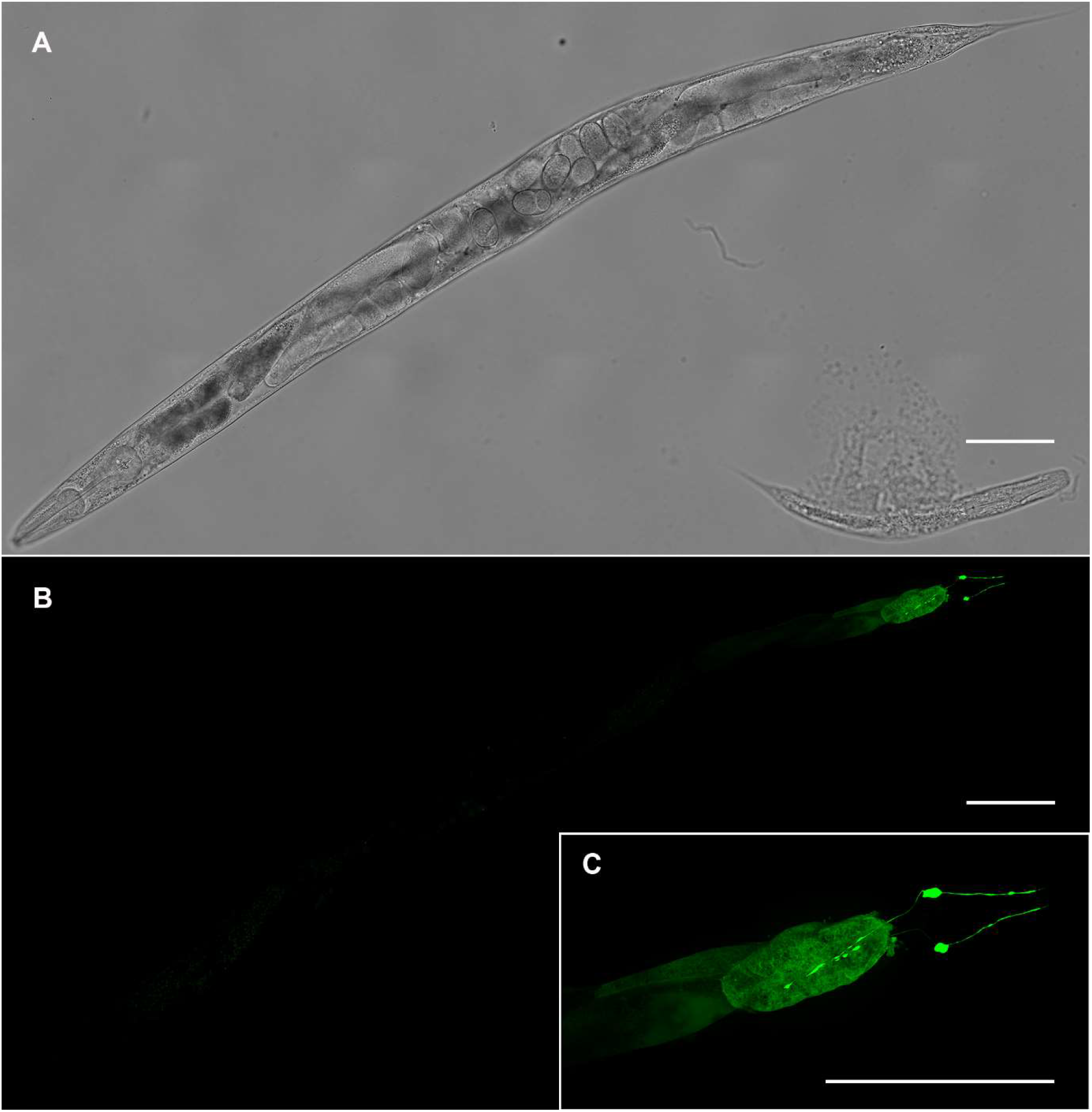
Localization of *gcp-2*.*2p::GFP* expression in the phasmids neurons (PHAL/R and PHBL/R) and their processes. A Bright-field image of adult worm. B The GFP signal was exclusively observed in phasmids neurons (PHAL/R) and their processes in the tail of adult *C. elegans*. C GFP signal in PHAL/R and PHBL/R phasmids in detail. Scale bars represent 100 µm.

**Figure 6.**
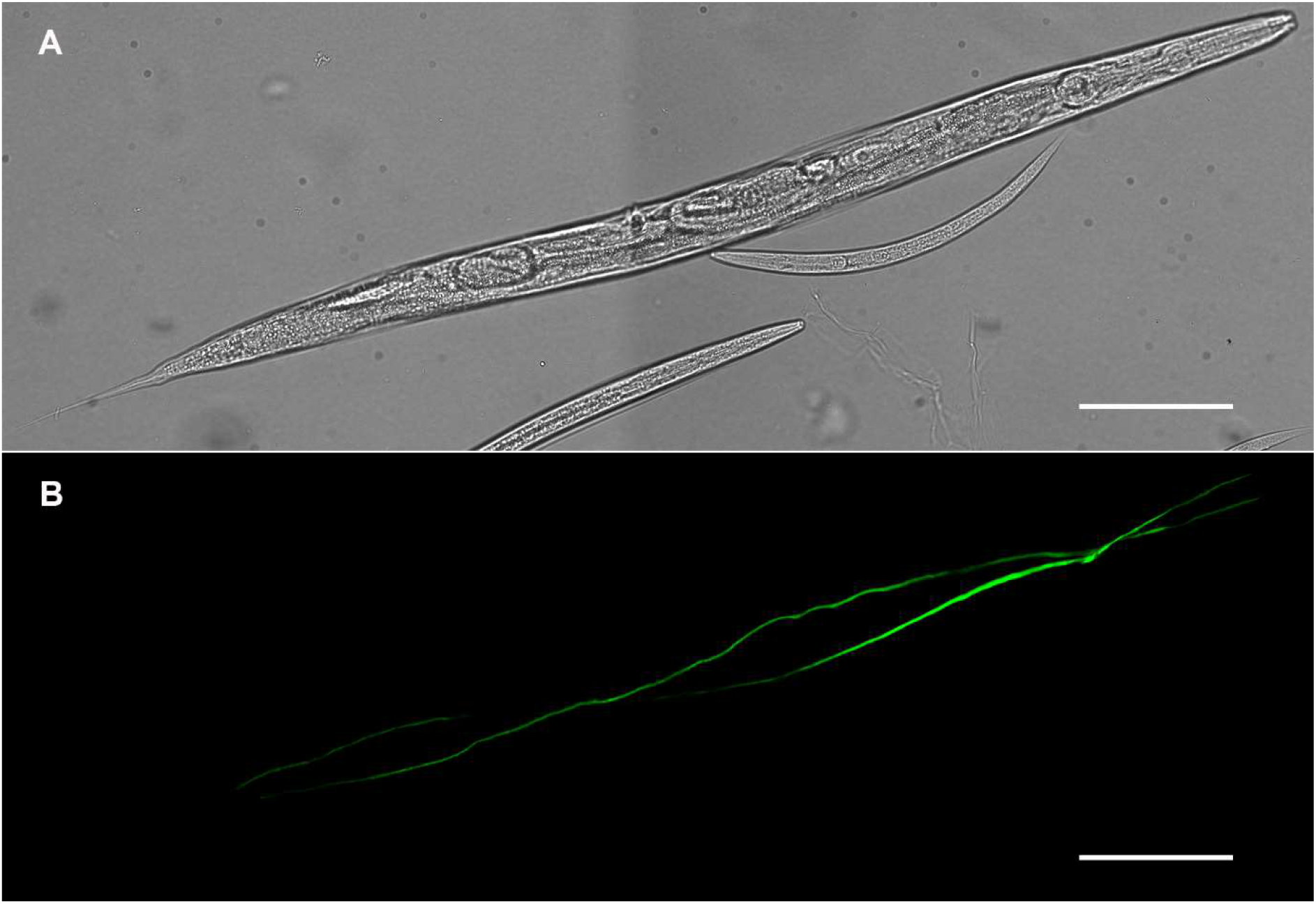
Expression of *gcp-2*.*3p::GFP* in the excretory cell of adult *C. elegans*. A Bright-field image of adult worm. B GFP expression pattern revealed the exclusive localization of *gcp2*.*3* in the excretory system (H-shape cell and its processes along the body of the worm). Scale bars represent 100 µm.

PHA and PHB, glutamatergic ciliated sensory neurons, are localized in the lumbar ganglia, exposed to the outside environment, and serve as chemosensory cells. Dye-filling with DiI was used for PHAL/R and PHBL/R neurons identification (Fig. EV4 B, C, D) and confirmation of phasmid’s functionality in *gcp-2*.*2* mutants (Fig. EV4 F). DiI is a lipophilic fluorescent dye able to selectively incorporate into two pairs of ciliated phasmid neurons in the tail (PHA and PHB) *via* cilia present at the tips of dendrites (Starich *et al*, 1995; Perkins *et al*, 1986).

### Knockout of peptidase paralogs is manifested in different phenotypes

Knockouts of the studied genes resulted in the most apparent phenotypes reported in any experimental animal to date.

The progeny production was changed after the *gcp-2*.*1* knockout. The number of progeny in the *gcp-2*.*1* mutant strain was reduced by 25% (Fig. 7 A; Appendix Table S1) compared to the wild-type strain but the difference was not statistically significant. Furthermore, manual counting revealed a 50% increase in pharyngeal pumping frequencies in the *gcp-2*.*1* mutant strain (Fig. 7 B; Appendix Table S2). At the same time, *gcp-2*.*1* knockout did not visibly affect the behavior and movement of the worms.

**Figure 7.**
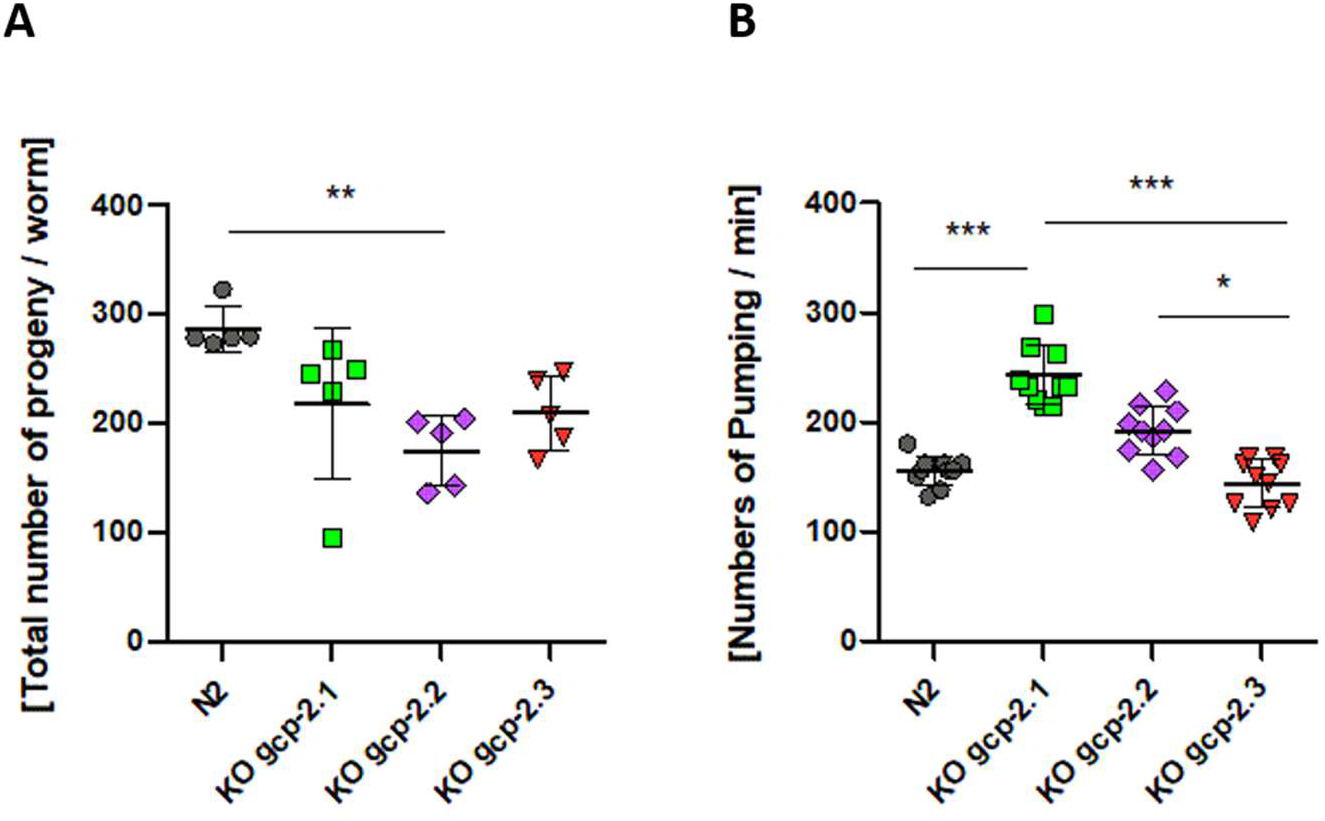
Effect of the gene knockouts on the phenotype manifestation. A Impact of gene knockouts on the reproduction of the worm. The number of offspring produced by one hermaphrodite mother of N2 wild-type strain and mutant strains *gcp-2*.*1, gcp-2*.*2, gcp-2*.*3* mutant worms. The statistical significance was determined by the Kruskal-Wallis test (nonparametric ANOVA, P=0.0054) and Dunn’s Multiple Comparison Test (Appendix Table S1). B Impact of gene knockouts on the pharyngeal pumping. Manual counting of pharyngeal pumping revealed the highest and the lowest rate of pumps for the *gcp-2*.*1* and *gcp-2*.*3* mutant strains, respectively. The data were tested by the Kruskal-Wallis test (nonparametric ANOVA, P < 0.0001) and Dunn’s Multiple Comparison Test (Appendix Table S2). P value < 0.001 - Extremely significant (***); P value 0.001 to 0.01 - Very significant (**); P value 0.01 to 0.05 – Significant (*).

Knockout of the *gcp-2*.*2* gene significantly negatively impacted progeny production rate (60% decrease compared to N2; Fig. 7 A; Appendix Table S1) and caused one day shorter progeny production. The increase in the pharyngeal pumping rate was not statistically significant (Fig. 7 B; Appendix Table S2). Interestingly, while handling the *gcp-2*.*2* knockout worms, we noticed they were more fragile compared to other mutants and wild-type worms. Thus, we performed a detailed ultrastructural analysis of the cuticle of all mutant strains using transmission electron microscopy (TEM). Figure 8 shows spongy formations in the basal zone of the cuticle, which were observed only in *gcp-2*.*2* mutant worms (Fig. 8 C and Fig. EV5).

**Figure 8.**
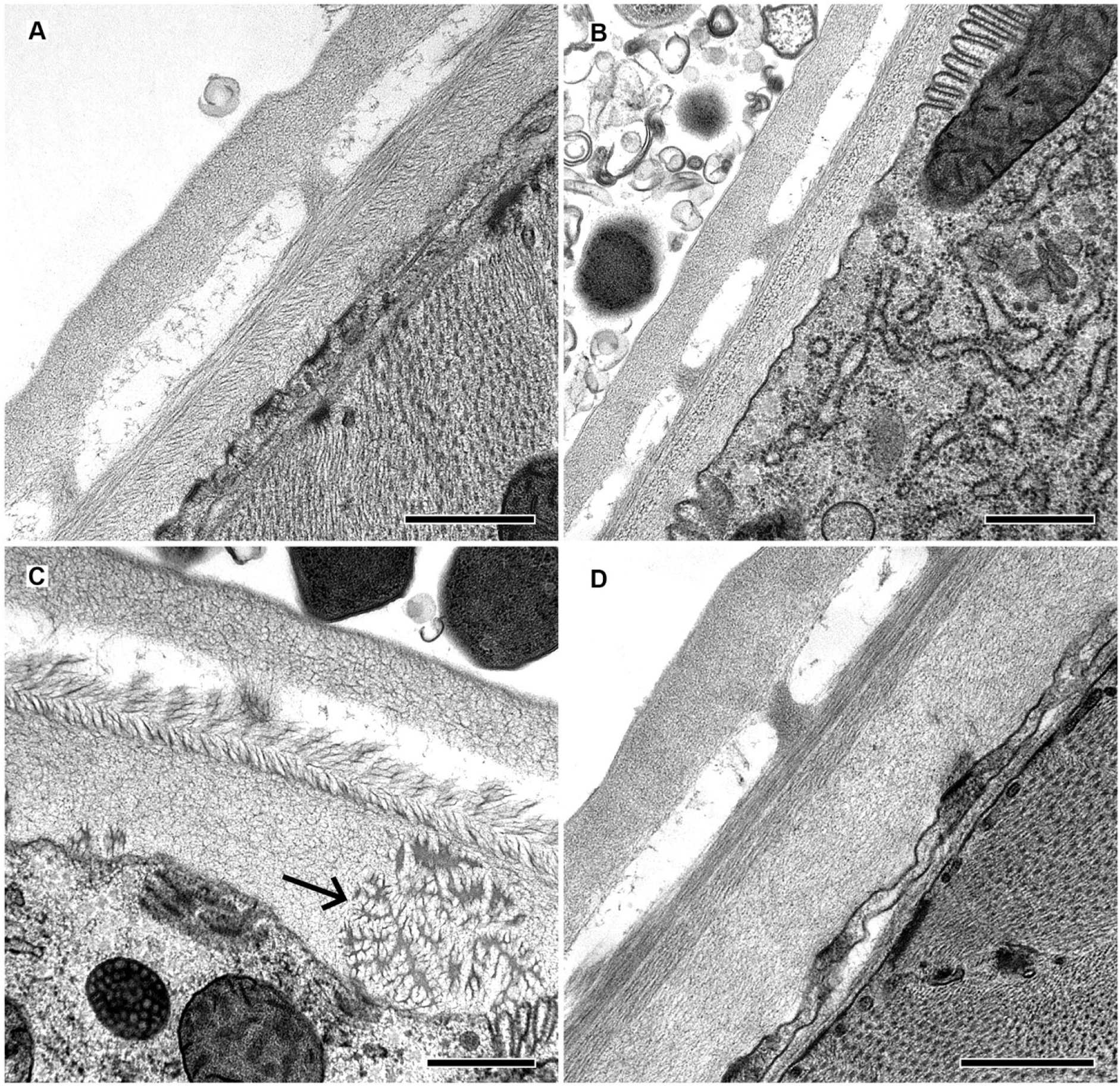
Ultrastructural analysis of the surface structures of *C. elegans*. A The image of cuticle of N2 strain. B Cuticle of *gcp-2*.*1* mutant adult worm. C Cuticle of *gcp-2*.*2* mutant adult worm. For *gcp-2*.*2* mutants (C) unusual spongy structures (arrow) in the basal layer were observed. These structures were not detected in the cuticle of other strains of *C. elegans* included in this study. D Cuticle of *gcp-2*.*3* mutant adult worm. Scale bars indicate 500 nm.

**Figure 9.**
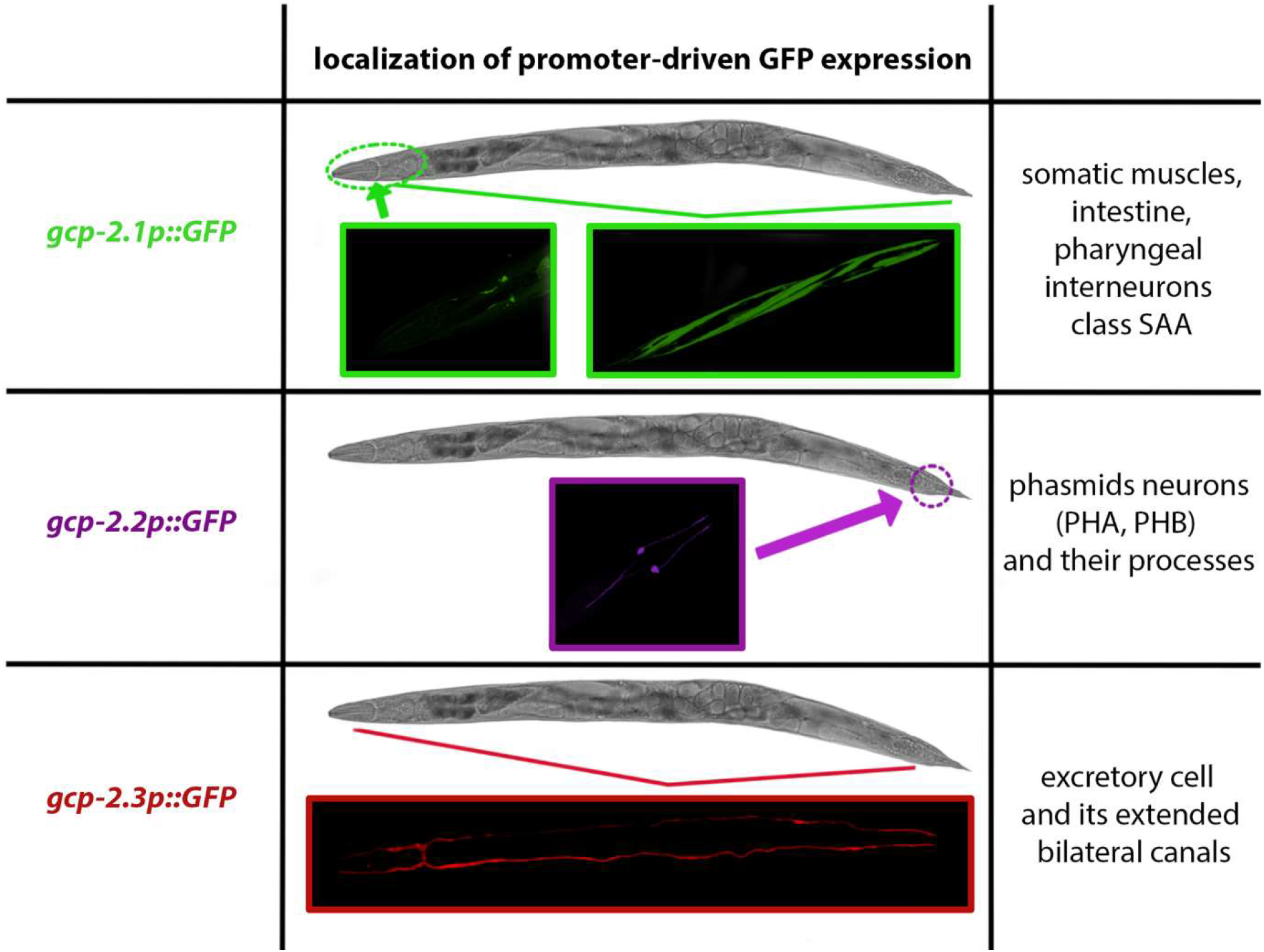
Overview of the promoter-driven GFP expression of three GCP2 paralogs in distinct tissues of *C. elegans*.

For *gcp-2*.*3* knockout worms, the number of offspring was reduced to approximately 75% of wild-type N2 controls, but the reduction was not statistically significant. (Fig. 7 A; Appendix Table S1). On the other hand, the pharyngeal pumping rate remained virtually unchanged and compared to that of N2 worms (Fig. 7 B; Appendix Table S2). The significant effect of the gene knockout on the lifespan of the mutants was not demonstrated (Appendix Fig. S2) (for overview see Table 2).

**Table 2:** Summary table of the most prominent phenotypes. Statistically significant changes in phenotype manifestation are marked in bold.

| phenotype | GCP2.1 | GCP2.2 | GCP2.3 |
| --- | --- | --- | --- |
| **pharyngeal pumping** | the highest increase in<br>the number of<br>pharyngeal pumping/s | increased rate of<br>pharyngeal pumping | lowest pharyngeal<br>pumping rate |
| **brood size** | reduced by a quarter | the lowest progeny<br>production | reduced by a quarter |
| **other observations** |  | pronounced fragility<br>spongy structure in the<br>cuticle |  |

## Discussion

The studies of the metalloproteases from the subfamily M28B primarily focus on the human GCP2 but significant gaps in the knowledge of the physiological functions of this protease and related orthologs still exist. By use of promoter-driven GFP expression together with gene knockout mutants, we were able to reveal some of the functions of these enzymes in *C. elegans* including unique phenotypes linked to the particular M28B metalloproteases in *C. elegans* (Bacich *et al*, 2002; Gao *et al*, 2015). Generally, observed phenotypic manifestations together with localizations indicate that the physiological functions of these enzymes may correspond to the already known functions and localizations of M28B metalloproteases in mammals.

Phylogenetic comparison of orthologs from the genus *Caenorhabditis* showed that CeGCP2.1 is phylogenetically relatively distant from CeGCP2.2 and CeGCP2.3, which are closely related to each other. The ancestor of CeGCP2.2 and CeGCP2.3 originated in gene duplication derived from CeGCP2.1, probably in the last common ancestor of the group *Caenorhabditidae*. Another consecutive gene duplication led to the speciation of CeGCP2.2 and CeGCP2.3, and there is strong evidence that these genes arose early in the evolution of the entire genus *Caenorhabditis*. They are likely more specialized for certain, yet unknown, physiological functions judging from their expression levels and localizations. Such duplication events occurred in nematodes several times independently, and still, many species including important parasites possess only a single gene (Howe *et al*, 2017), making those enzymes putative drug targets.

Homology modeling showed that residues coordinating Zn^2+^ ions and the proton shuttle Glu424 essential for catalytic activity are conserved in all *C. elegans* GCP2 orthologs. Human GCP2 exhibits a preference for negatively charged substrates containing glutamate (Barinka *et al*, 2002), while some changes in structural elements of *C. elegans* orthologs indicate possible differences in substrate preference. Residues that bridge zinc ions in the hsGCP2 structure (Asp387, His377, Asp453, Glu425, His553) (Mesters *et al*, 2006) are conserved in all three *C. elegans* orthologs; this suggests that the overall structure described for human GCP2 is the same for *C. elegans*. Residues Arg210, Tyr552, Tyr700, Asn257 and Lys699 are crucial for substrate recognition for hsGCP2 (Mesters *et al*, 2006). Those residues are also conserved in all three *C. elegans* orthologs, whereas Asn257 is replaced in all three *C. elegans* orthologs by serine, which is also a polar amino acid with an uncharged side chain. This indicates that the structural function remains conserved. Other critical residues of hsGCP2 for substrate residue recognition are the S1 pocket residues - Asn519, Arg463, Arg534, Arg563 (Barinka *et al*, 2008). Lys699 is replaced by serine in CeGCP2.1 and by tryptophan in CeGCP2.2 and CeGCP2.3. This means significant changes: replacement of a positively charged amino acid by a polar uncharged serine, and by an amino acid with a hydrophobic side, in CeGCP2.1 and CeGCP2.2 together with CeGCP2.3, respectively. In the S1 pocket of hsGCP2, Tyr549, and Tyr552 are important hydrophobic residues. Tyr 549 is only substituted for isoleucine in GCP2.3, so even after this substitution the hydrophobicity and thus the structural function is preserved.

The protein alignment of the primary structures of all three studied proteases and hsGCP2 reveals, that CeGCP2.3 lacks the transmembrane domain and has a partial deletion of the C-terminal dimerization domain. This suggests that the CeGCP2.3 protein might be intracellular or potentially even secreted (Jeffery, 2020; Jang *et al*, 2002; Mazumder *et al*, 2010). The results of the protein structure analysis showed the closest similarity between CeGCP2.2 and CeGCP2.3, while CeGCP2.2 and CeGCP2.1 are less similar. Also, CeGCP2.1 is more similar to its human ortholog hsGCP2 than to the *C. elegans* paralog CeGCP2.3 (Table 1).

Although *gcp-2*.*3* is annotated as a pseudogene in the current version of WormBase (WS284) (Davis *et al*, 2022), our RT-qPCR data and transcriptomic data (unpublished) together with previously published time-resolved (Boeck *et al*, 2016) and tissue-specific transcriptomes (Kaletsky *et al*, 2018) of *C. elegans* showed that *gcp-2*.*3* is indeed transcribed. Time-resolved transcriptome of *C. elegans* showed that *gcp-2*.*3* is expressed mainly in embryonal stages where it reaches its peak expression 0.6 DCPM (depth of coverage per million reads) dauer larvae where it reaches up to approximately 0.45 DPCM. In other stages expression is much lower and is bellow 0.2 DPCM (Boeck *et al*, 2016).

Transcriptomic data for L2 tissues (Boeck *et al*, 2016) showed that *gcp-2*.*2* is expressed in pharyngeal muscles, ciliated sensory neurons, excretory cells, non-seam hypodermis, body wall muscles, and intestine, while expression of *gcp-2*.*3* was reported in socket cells and intestines. In addition, tissue-specific transcriptomic analysis of adults (Kaletsky *et al*, 2018) showed that *gcp-2*.*2* is mainly expressed in pharyngeal gland cells, RID (single motor neuron, interneuron situated in the dorsal ganglion, innervates dorsal body muscles of *C. elegans*), inner labial neurons, head muscles, GABAergic neurons, head neurons, rectal gland cells, and head ganglion. For *gcp-2*.*3*, the expression is limited to amphid sheath cells, RID, and phasmid neurons. Contrary to these findings, our promoter-driven GFP reporters revealed different localizations. *Gcp-2*.*2p::GFP* expression was restricted to phasmids (PHA and PHB) in both larvae and adult worms and *gcp-2*.*3p::GFP* was exclusively expressed in the excretory cell and its processes of larvae and adult wild-type worms. Discrepancies between the results could be because the above-mentioned transcriptomic datasets served primarily as expression prediction, and they do not provide information about the spatial constraints of expression. Moreover, our design of the GFP-fusion construct for *gcp-2*.*2* and *gcp-2*.*3* may not cover all regulatory elements in the different tissues and cells.

Based on our localization studies, *gcp-2*.*1* is expressed in pharyngeal interneurons class SAA, somatic muscles, and intestine. SAA class neurons belong to neurons receiving cholinergic inputs (Pereira *et al*, 2015). The long, anteriorly directed processes of SAA interneurons probably serve as stretch receptors monitoring the posture of the tip of the head (White *et al*, 1986). SAA interneurons interact with the body and the head motor system, and it seems that they can coordinate head and body movement (White *et al*, 1986). Indeed, localization of *gcp-2*.*1* expression in muscles, nervous system, and SAA interneurons could be related to some of our phenotyping results. We found out that *gcp-2*.*1* mutant worms showed the most significant increase in the number of pharyngeal pumps per minute of all studied paralogs. Even if the pumping rate is modulated by multiple mechanisms (Trojanowski *et al*, 2016), the elimination of the *gcp-2*.*1* gene has an impact on the speed of pumping. It could suggest that CeGCP2.1 plays an important role in pharyngeal pumping modulation. Although the somatic muscles, nervous system, and intestine were dominant locations of the *gcp-2*.*1* expression, we did not detect any other phenotypic manifestations related to the digestive or locomotor system in *gcp-2*.*1* mutant worms. It is also possible that the expression in the intestine is non-specific (Hunt-Newbury *et al*, 2007). Despite the abnormal pharyngeal pumping rate, which could have an impact on the fitness of the worms, knockout of the *gcp-2*.*1* had only a mild effect on fertility. Knockout of the *gcp-2*.*2* gene caused increased fragility of *C. elegans* cuticle. This unique fragile phenotype was accompanied by previously undescribed spongy-structure formations in the basal zone of the cuticle which we presume to be responsible for the observed fragility of worms during the routine manipulation. We also observed the lowest offspring production of all compared strains. Theoretically, cuticle fragility could affect offspring production by weakening the vulval cuticle, but we did not detect bursting through the vulva or other indicators of defective egg laying; therefore, these two phenotypes (lower fecundity and brittle cuticle) are likely unrelated. However, the localization of *gcp-2*.*2* expression pointing to phasmid neurons (PHA, PHB) does not explain described fragility and reduction in offspring production. The reason may be that we were not able to detect all regulatory elements for epidermal expression and thus did not localize the gene expression in the cuticle.

The expression of *gcp-2*.*3* was localized to the excretory cell, which maintains osmotic equilibrium in the worm body (Sundaram & Buechner, 2016), and its extended bilateral canals reaching from the nose of the worm to the tail region. Many mutations that affect the excretory system cause a lethal phenotype, or mutants with milder abnormalities of these cells often appear pale, and slightly bloated (Sundaram & Buechner, 2016). Nevertheless, such phenotypes were not noticed in the case of *gcp-2*.*3* mutant worms during our study. Knockout of the *gcp-2*.*3* led to a decrease in fertility similarly as observed in the case of the *gcp-2*.*1* mutant worms. We are not able to state that the lower fertility is directly affected by the missing expression of *gcp-2*.*3* in the excretory cell. But knockout of the *gcp-2*.*3* might negatively influence the fitness of the animals and thus led to a decrease in progeny.

Concerning the phenotype manifestation of *C. elegans gcp-2* gene knockouts and their particular localizations, there are noticeable significant similarities with mammalian protease orthologs. Among others, these proteases have been very well-described and studied in the digestive system (Rais *et al*, 2016; Visentin *et al*, 2014; Pangalos *et al*, 1999) and also in the brain, astrocytes, and Schwann cells (Rovenska *et al*, 2008). Similarly, our observations revealed the expression of *C. elegans gcp-2*.*1* in the intestine and nervous tissue of adult worms. However, our data from *in silico* modeling and a recent experiment with the recombinant protease (not published) do not support the possibility that CeGCP2.1 would copy the physiological function of human GCP2 by removing terminal glutamate.

Using the transcript information from WormBase (Harris *et al*, 2020), we analyzed two additional CeGCP2.1 splice variants. A comparison of CeGCP2.1 isoforms revealed that isoform CeGCP2.1b lacks the predicted intracellular domain. Isoform CeGCP2.1c completely lacks the C-terminal domain, which implies the loss or significant changes in the catalytic function, given that in its human homolog GCP2, the C-terminal domain is involved in substrate recognition (Barinka *et al*, 2007; Mesters & Hilgenfeld, 2008). No biological functions associated with these isoforms of GCP2.1 protein have been described so far, so we can only conclude homology to other isoforms and GCP2-like proteins of *Caenorhabditis* species, NAAG and transferrin receptors.

We have demonstrated that the *gcp-2*.*2* mutant line almost completely suppressed the expression of the *gcp-2*.*3* gene; the decreased expression could be explained by the aforementioned proximity of both genes on the chromosome, suggesting the presence of an alternative polycistronic expression of these proteins (Evans & Blumenthal, 2000). Additionally, we have found that knockout of the *gcp-2*.*1* gene significantly increased the expression of *gcp-2*.*3*, which is surprising given the differences in phenotypic expression and localization. The question is whether the increased expression of the *gcp-2*.*3* gene can somehow compensate for the missing *gcp-2*.*1*. In addition, the knockout of the *gcp-2*.*3* gene resulted in a decrease in *gcp-2*.*1* and *gcp-2*.*2* expression. A possible explanation for this decrease in *gcp-2*.*1* expression could be impaired function of the excretory system and thus a decrease in the fitness after the *gcp-2*.*3* gene knockout. However, we do not yet have a satisfactory explanation for these changes in expression.

To summarize, we have used *C. elegans* as a tool to describe the diversification of one gene; on a similar principle, we can hypothesize an indirect parallel of the diversification of five independently originated genes in humans. Based on localization and phenotyping studies, we have shown that *gcp-2*.*3* is not a pseudogene, as inaccurately annotated in the WormBase database (Harris *et al*, 2020), but an active paralogous gene. Precise localization using promoter-driven GFP expression revealed strict expression patterns for each of three *C. elegans* genes. The localization of these *C. elegans* proteases from the M28B family is strikingly analogous to their mammalian orthologs. The knockout of the genes studied revealed the strongest phenotypic manifestations ever observed in the animal kingdom. In addition to the impact on pharyngeal pumping and reproduction of the mutant worms, we described a unique phenotype of the fragile worm, probably caused by abnormalities (spongy formations) in the worm cuticle.

## Materials and methods

### Sequence alignment and homology modeling

The amino acid sequence of hsGCPII (NCBI ID: NP_004467.1), and *C. elegans* orthologs deposited in the UniProtKB: GCP2.1 under sequence ID: P91406-1 coded by *gcp-2*.*1* (R57.1) gene; GCP2.2; under sequence ID: Q93332 coded by *gcp-2*.*2* (C35C5.2) gene; and GCP2.3 translated from the sequence inaccurately labeled as a pseudogene *gcp-2*.*3* (C35C5.11) from WormBase under sequence ID: C35C5.11, were used for the sequence alignment and homology modeling. The amino acid sequences of hsGCPII and all three *C. elegans* orthologs were aligned using Jalview Version 2 (Waterhouse *et al*, 2009) and subsequently analyzed for sequence identity and similarity with Sequence Manipulation Suite: Ident and Sim tool (Stothard, 2000).

For homology modeling, the 5ELY GCP2 structure determined to high (1.81 Å) resolution limits was selected as a template (Novakova *et al*, 2016). The Modeller 9.23 software was used to construct the target-template sequence alignment and to generate five 3D homology models. The best model for each enzyme was selected based on discrete optimized protein energy (DOPE) scores (Webb & Sali, 2016). Finally, the homology models of CeGCP2 variants and hsGCPII structure (PDB code 3BXM) were superimposed in PyMol and analyzed by visual inspection.

The amino acid sequences of hsGCPII and all three *C. elegans* orthologs were aligned using Jalview Version 2 (Waterhouse *et al*, 2009) and subsequently analyzed for sequence identity and similarity with Sequence Manipulation Suite: Ident and Sim tool (Stothard, 2000).

### Phylogenetic analysis

For phylogenetic analysis, protein sequences of nematode paralogs of the M28B peptidase family were downloaded from the MEROPS database (Rawlings *et al*, 2018) and aligned with sequences acquired from NCBI and WormBase databases (Davis *et al*, 2022; Sayers *et al*, 2022). The alignment was constructed by MAFFT v. 7.222 and automatically trimmed by BMGE v. 1.12 software using the blosum62 scoring matrix. The phylogenetic tree was constructed in IQ-TREE v. 1.6.1 using the best fitting model (LG4M) selected by ModelFinder. GCP2 of *Homo sapiens* was used as the outgroup. The resulting tree was graphically enhanced in Figtree v. 1.4.4 (http://tree.bio.ed.ac.uk/software/figtree).

### Maintenance of *C. elegans* and backcrossing

All *C. elegans* nematodes were grown and maintained for several generations at 18 °C under standard conditions (Stiernagle, 2006). Worms were kept on nematode growth medium (NGM) plates and fed by OP50 *E. coli* strain, which was performed according to slightly modified instructions in the WormBook (Stiernagle, 2006). Three *gcp-2* mutant worm strains RB1055 - gcp-2.1 (ok1004) (CGC), gcp-2.2 - gcp-2.2 (tm6541) and C35C5.11 - gcp-2.3 (tm5414) (MITANI, 2009), and the wild-type (N2) were used in this study (Appendix Table S3). The mutation of the three paralog genes represents loss-of-function mutation.

Before all experimental procedures, knockout worms were backcrossed eight times with N2 males with the consequent recovery of mutant homozygotes to remove possible off-target and other random genetic mutations. Backcrossing was performed according to WormBook protocols (Ahringer, 2006).

DNA for control genotyping was isolated from a single hermaphrodite after the offspring production. The worms were dissolved in 10ul of QuickExtract(tm) DNA Extraction Solution (Biosearch Technologies). Final analyses of homozygous or heterozygous lines were performed by PCR reaction using Ready To Use PCR MasterMix (Central European Biosystems) with forward and reverse primer for the target gene (Appendix Table S4), and resolved by agarose gel electrophoresis (see Appendix Table S5, Appendix Fig. S3 and detailed protocols therein).

### Gene expression assay

Adult worms were washed down from the growing plate with PBS buffer (Sigma Aldrich) and spun down at 3000 g. Worms were resuspended in 500 µl of TRIzol(tm) Reagent (ThermoFisher Scientific) and RNA was isolated with RNeasy Mini Kit (Qiagen) according to manufacturer instructions. Any remaining DNA was removed by TURBO(tm) DNase (Thermo Fisher Scientific). The reverse transcription to obtain cDNA from RNA samples was performed with SuperScript(tm) III (Thermo Fisher Scientific) and Oligo(dT)18 primers according to the manufacturer’s protocol. Resulting cDNA samples were stored at – 20 °C until use.

The RT-qPCR analysis of all three paralog genes (*gcp-2*.*1, gcp-2*.*2, gcp-2*.*3*) and a housekeeping gene *C. elegans tubulin* alpha (TBA1, (Hoogewijs *et al*, 2008)) was performed for each sample in biological triplicates by use of LightCycler® 480 SYBR Green I Master mix (Roche) and evaluated by LightCycler® 480 II cycler (Roche) according to the manufacturer’s instructions. Gene-specific primers were selected and evaluated as previously published (Fajtová *et al*, 2015).

### Behavioral phenotyping

All behavioral observations of knockout worms were carried out at controlled room temperature (21 - 22°C). To determine the development speed, worms were age-synchronized according to WormBook protocol (Stiernagle, 2006). Then, synchronized L1 stages were seeded on the NGM plates under standard cultivation conditions and monitored until they reached the young adult stage. Age-synchronized young adults were placed on individual plates each day until they ceased reproducing. Fecundity was measured by manual counting of viable progeny and unviable embryos (Maulik *et al*, 2017). Feeding behavior was visually scored by counting the number of pharyngeal pumps. A single pharyngeal stroke was defined as one synchronous contraction and relaxation cycle of the corpus and terminal bulb (Raizen *et al*, 2012). The worms (L4-stage) were placed on plates under a stereomicroscope Nikon SMZ 25 and recorded for 15 s using a Nikon DS-Ri2 camera. Then the pumping rate was quantified for 10 worms from each strain using a hand-held counter in 15 s frames 10 times for each animal. The mean and standard deviation were counted. The experiment was repeated three times with the same results.

### Localization studies by GFP promoter-driven expressions

Promoter regions (entire upstream non-coding regions) of three studied genes were cloned into the plasmids optimized for *C. elegans* GFP expression. Upstream promoters areas for *gcp-2*.*1, gcp-2*.*2*, and *gcp-2*.*3* genes in sizes 2.893 bp, 1.107 bp, and 911 bp (Appendix Fig. S4) respectively were PCR amplified by specific primers (Table S2) using Phusion^®^ High-Fidelity PCR Master Mix (Thermo Fisher Scientific) from genomic DNA template and sub-cloned by restriction cloning into GFP expression plasmids pPD95_79 and pPD95_81. pPD95_79 and pPD95_81 kindly provided by Andrew Fire (pPD95_79 - Addgene plasmid #1496; http://n2t.net/addgene:1496; RRID:Addgene_1496and pPD95_81 - Addgene plasmid #1497; http://n2t.net/addgene:1497; RRID:Addgene_1497).

The resulting isolated plasmids were used for microinjections as described previously (Evans, 2006). Used injection mix was composed of GFP-expression plasmid (10 ng/μl), pRF4:rol-6(su1006) co-injection marker (50 ng/μl) (Mello *et al*, 1991), pBluescript (Addgene) empty plasmid (180 ng/μl) was added to reach a total DNA concentration of 100-200 ng/μl in sterile ddH2O. Agarose pads (2%) and Halocarbon oil 700 (Sigma-Aldrich) were used for imobilization, and a standard M9 medium was used as a recovery buffer (Evans, 2006). Worms phenotype (Rol of animals) for control co-injection marker (pRF4:rol-6(su1006)) was checked under the microscope. Transformed worms were picked and put on a new seeded plate. Transgenic lines were then maintained according to standard protocols (as described above).

### In vivo analysis and imaging of tissue-specific GFP expression in C. elegans

Transgenic worms were anesthetized using 20mM sodium azide (NaN3) in M9 medium, placed on a 5% agarose pad on glass slides, and gently covered with cover slides. All images were acquired using a spinning disc confocal microscope (Nikon CSU-W1) using a 60x WI objective (CF Plan Apo VC 60XC WI). Representative images are shown as a projection of X-Y z-stacks using the maximum intensity projection type. Images were analyzed using Huygens Professional 19.10 and open-source ImageJ software (Schneider *et al*, 2012).

### Staining of phasmids by DiI

DiI staining of phasmids was performed according to the protocol created by Michael Koelle and published on https://www.wormatlas.org/EMmethods/DiIDiO.htm. DiIC12(3) (1,1’-Didodecyl-3,3,3’,3’-Tetramethylindocarbocyanine Perchlorate) (Invitrogen, D383) was used. DiI is the lipophilic fluorescent dye, which can label phasmid neurons PHA and PHB (Tong & Bürglin, 2010).

### Electron microscopy

*C. elegans* worms, including *E. coli* OP50 bacteria, were scraped onto the high-pressure freezing (HPF) specimen carriers. Bacteria served as a filler, minimizing water content and facilitating freezing (Mulcahy *et al*, 2018). Worms on the carriers were frozen in the high-pressure freezer (Leica EM PACT2). Frozen samples collected on metal carriers were transferred under liquid nitrogen into a pre-frozen cryotube containing 1 ml freeze-substitution solution (2% OsO4 in 100% acetone with 1% lecithin) and finally to a freeze substitution unit (Leica EMAFS) for processing. Samples embedded in Epon EmBed812 resin were ultrasectioned (80nm) with an ultramicrotome (Leica EM UC 6) and placed on copper mesh 300 previously coated with uranyl acetate and lead citrate. Sections were finally examined using a transmission electron microscope (JEOL JEM 2100-Plus 200kV).

### Statistical analysis

Evaluation of the statistical data of mutant phenotypes (gene expression, fertility, pharyngeal pumping) was processed by GraphPad Prism 5.

## Supporting information

Expanded view

## Acknowledgement

We would like to acknowledge Luisa Cochella, Ph.D. for her very valuable and helpful advice and assistance in the field of transgenic approaches, transgenic worm screening, worm manipulation, and for the help and work she and her team at the Research Institute of Molecular Pathology (IMP, Vienna) spent on transgenic worm preparation. We thank to Dr. Lucie Jedlickova for and excellent graphic work. We also acknowledge Imaging Methods Core Facility at BIOCEV, institutional support by the MEYS CR (Large RI Project LM2018129 Czech-BioImaging) and ERDF (project No. CZ.02.1.01/0.0/0.0/18_046/0016045) for their support with obtaining imaging data presented in this paper.

JD and CB were supported by the Czech Science Foundation (Grant No. 18-14167S) and JD was supported the Czech Ministry of Education, Youth and Sports (Grant No. LTAUSA19023). The funding source(s) was not involved in the study design, the collection, analysis and interpretation of data, the writing of the report or in the decision to submit the article for publication. Infrastructural support was provided by the Czech Academy of Sciences (RVO: 86652036), and by the Czech University of Life Sciences, NutRisk Centre, (Grant No. CZ.02.1.01/0.0/0.0/16_019/0000845).

N2 and RB1055 strains were provided by the CGC, which is funded by the NIH Office of Research Infrastructure Programs (P40 OD010440). *gcp-2*.*2* and C35C5.11 strains were provided by the NBRP, which is funded by the Japanese government.

## Author contribution

JD, CB and MM were involved in conceptualization; JD, CB, MM, JS, MA participated in the methodology; LP, SN, DK, FK, ZK, LK, CB, MM and VV were involved in formal analysis; LP, SN, DK, FK, CB, JD were involved in the investigation; LP, SN, VV, MM, JD were involved in the data curation; LP, SN, JD were involved in writing - original draft preparation; VV, LK, ZK, CB, MA, JS, MM were involved in the draft review and editing; JD was involved in supervision and project administration. All authors have read and agreed to the published version of the manuscript.

