## Supplementary material for "Uncovering the essential roles of human GCP 2 orthologs in *Caenorhabditis elegans*": Expanded view

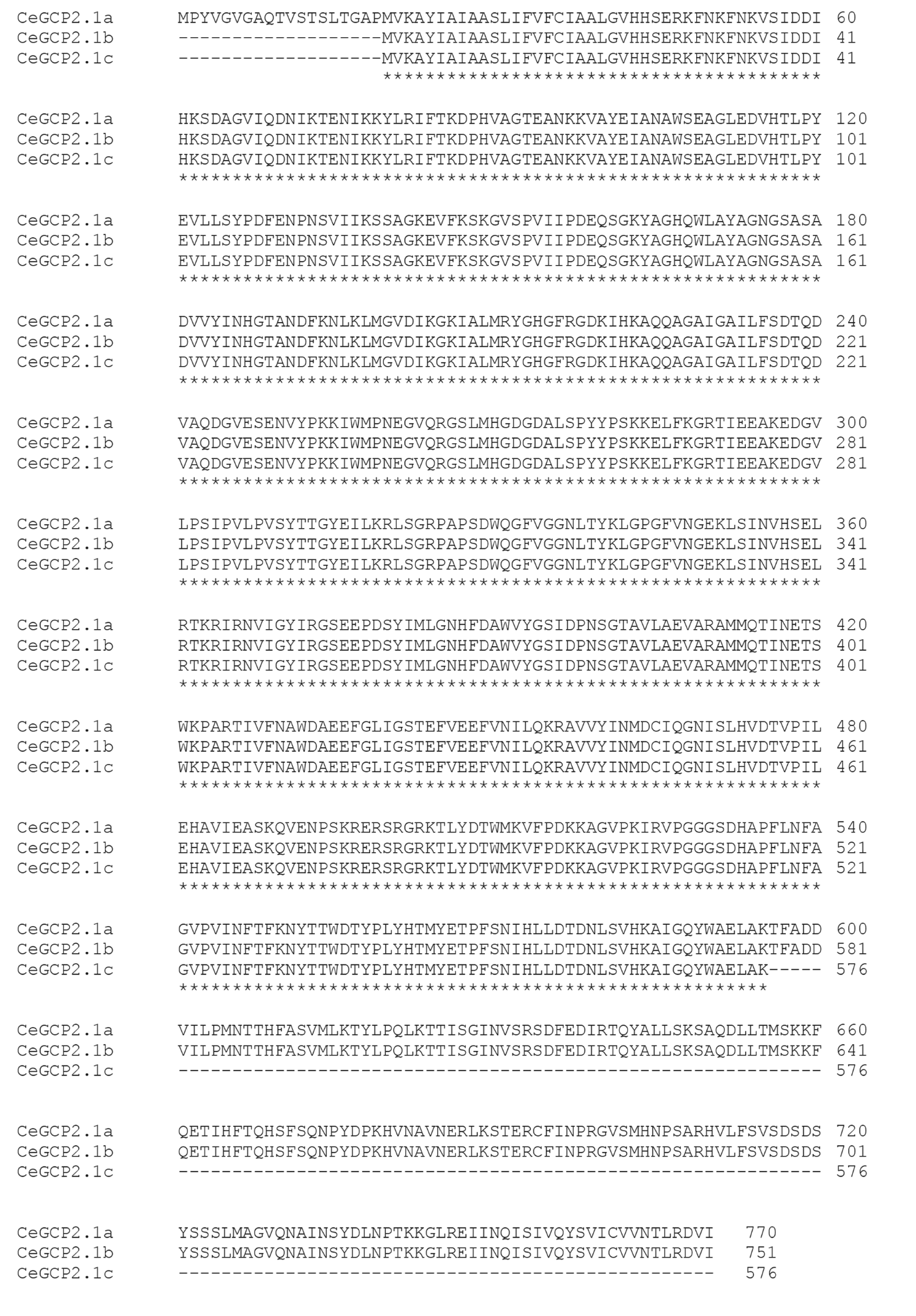

**Figure EV1 –** **Primary sequence alignment of three *C. elegans* GCP2.1 splice variants.** Performed using the Clustal Omega multiple protein alignment tool.(Sievers *et al*, 2011) Sequences were downloaded from the WormBase database (Davis *et al*, 2022) (WS285 version of WormBase) – GCP-2.1, isoform a (WormBase ID : R57.1a), GCP-2.1, isoform b (WormBase ID : R57.1b), and GCP-2.1, isoform c (WormBase ID : R57.1c).

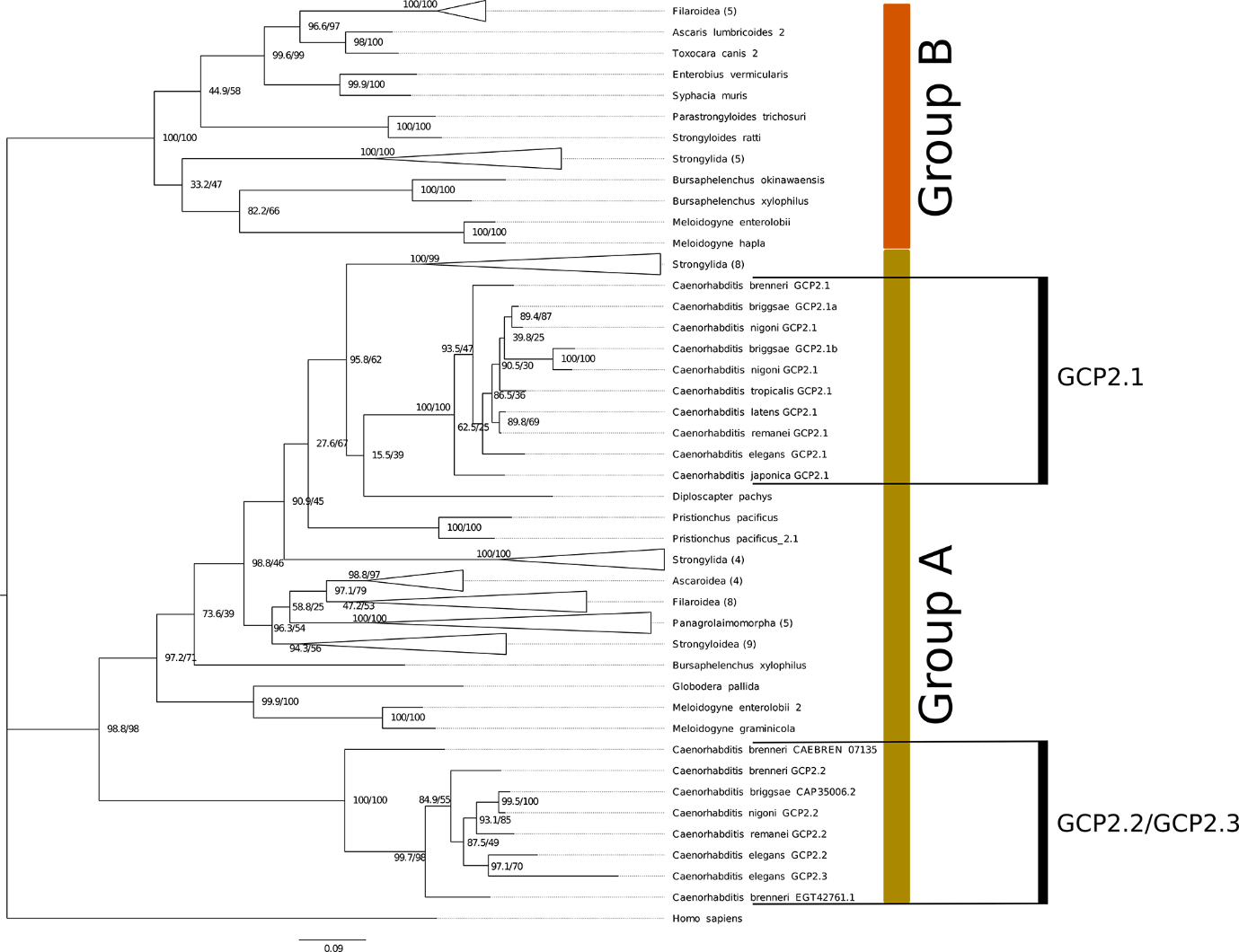

**Figure EV2 – Maximum likelihood phylogeny tree showing the relationship between M28B peptidases of nematodes.**

In the phylogeny tree *gcp-2.2* and *gcp-2.3* form a well-supported clade with their homologs from the genus *Caenorhabditis* showing that those genes evolved separately from *gcp-2.1*. The tree was constructed in IQ-TREE v 1.6.1 according to the best-fitting model (LG4M) and rooted by *Homo sapiens* GCP2. Numbers on nodes represent standard bootstrap and SH-aLRT support.

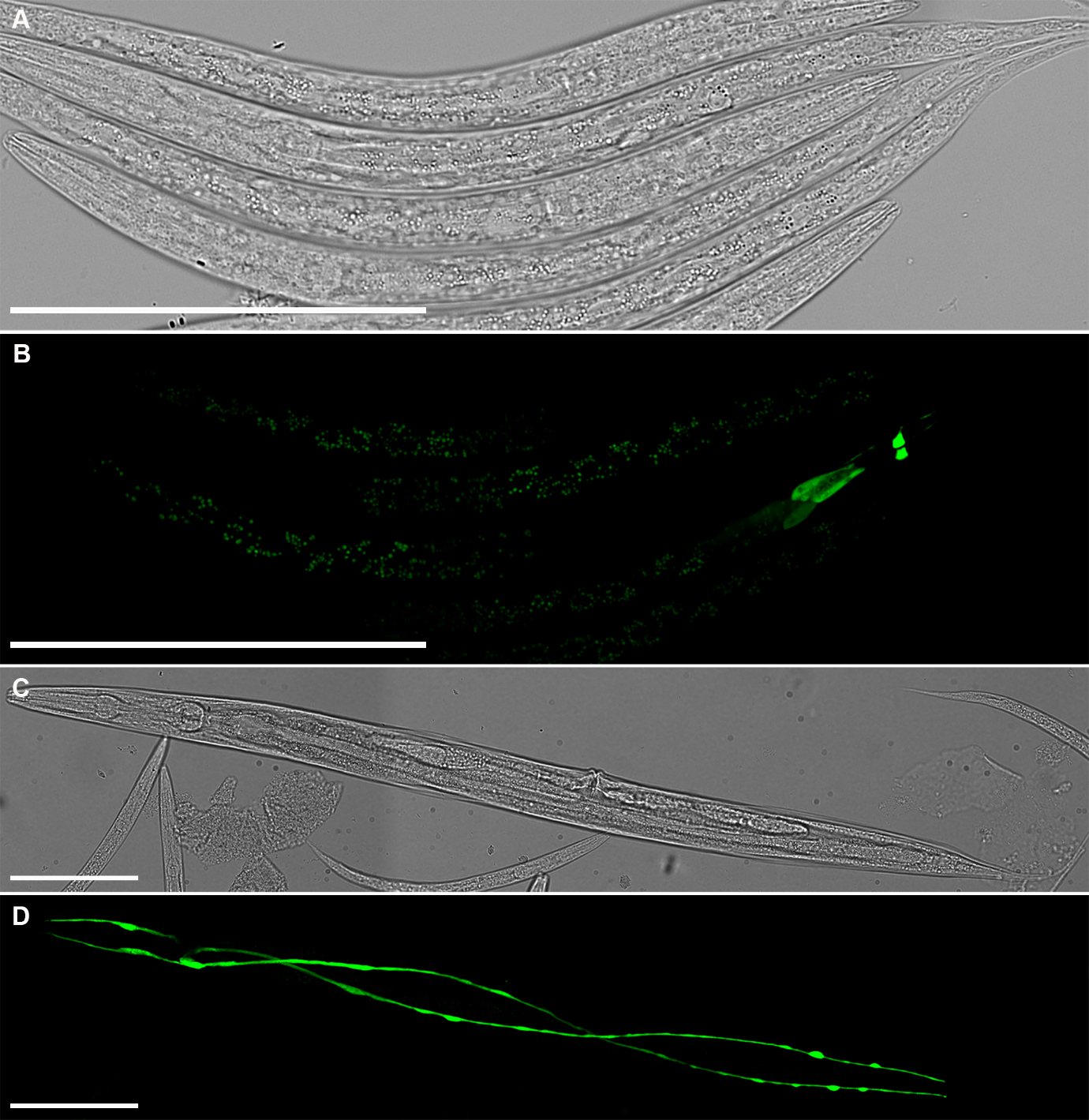

**Figure EV3 - Localization of *gcp-2.2p::GFP* expression in the phasmid neurons (PHA and PHB) and their processes of L3 larvae.**

A Bright-field image of *C. elegans* L3 larvae

B Same as for *gcp-2.2* in adults, the GFP signal corresponding to *gcp-2.2* expression was exclusively observed in phasmids neurons (PHAL/R and PHBL/R) and their processes in the tail of *C. elegans* larvae. One larva carrying GFP reporter construct among the negative larvae. The mosaicism is due to the extrachromosomal expression of the transgene.

**C, D Expression of *gcp-2.3p::GFP* in the excretory tissue of L4 larvae of *C*. *elegans*.**

C Bright-field of L4 larvae

D The localization of *gcp-2.3p::GFP* expression was the same as in adults, and it was exclusively located in the excretory canal cell (H-shape cell and its processes).

The scale bars represent 100 µm.

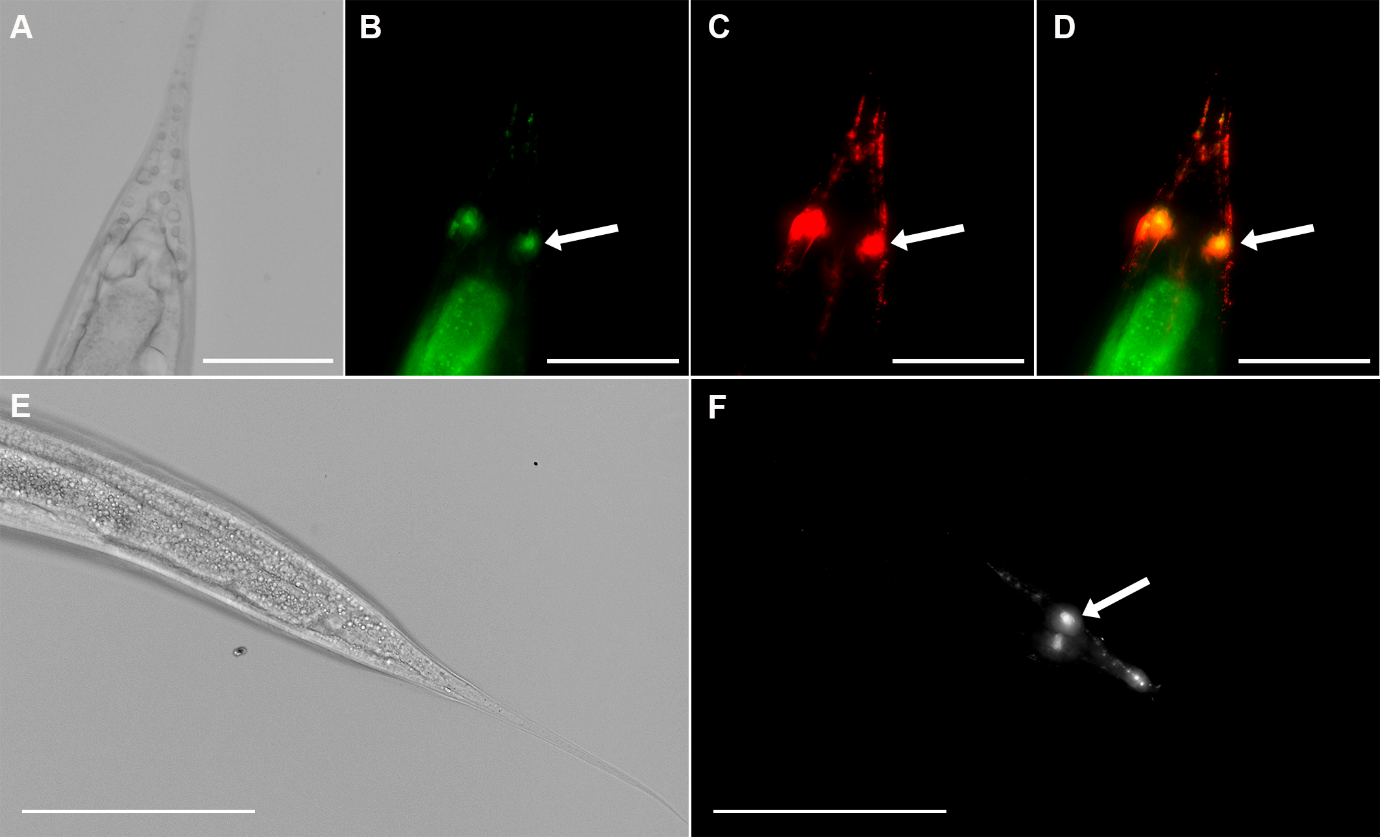

**Figure EV4 – Identification of a pair of phasmid neurons PHA and PHB in a transgenic *C. elegans* worm.**

A DIC image of *C. elegans* tail.

B Localization of *gcp-2.2p::GFP* expression.

C DiI stained phasmid neurons (PHAL/R, PHBL/R).

D Merged images allow to see that *gcp-2.2p::GFP* expression corresponds to DiI filling phasmid neurons PHA and PHB.

Arrows indicate phasmid neurons. Scale bars represent 50 µm.

E DIC image of *C. elegans* tail.

F Fluorescent visualization (DiI signal) of phasmid sensory neurons (arrow) and their processes in the tail of *C. elegans* *gcp-2.2* KO L4 larvae. Scale bars represent 100 µm.

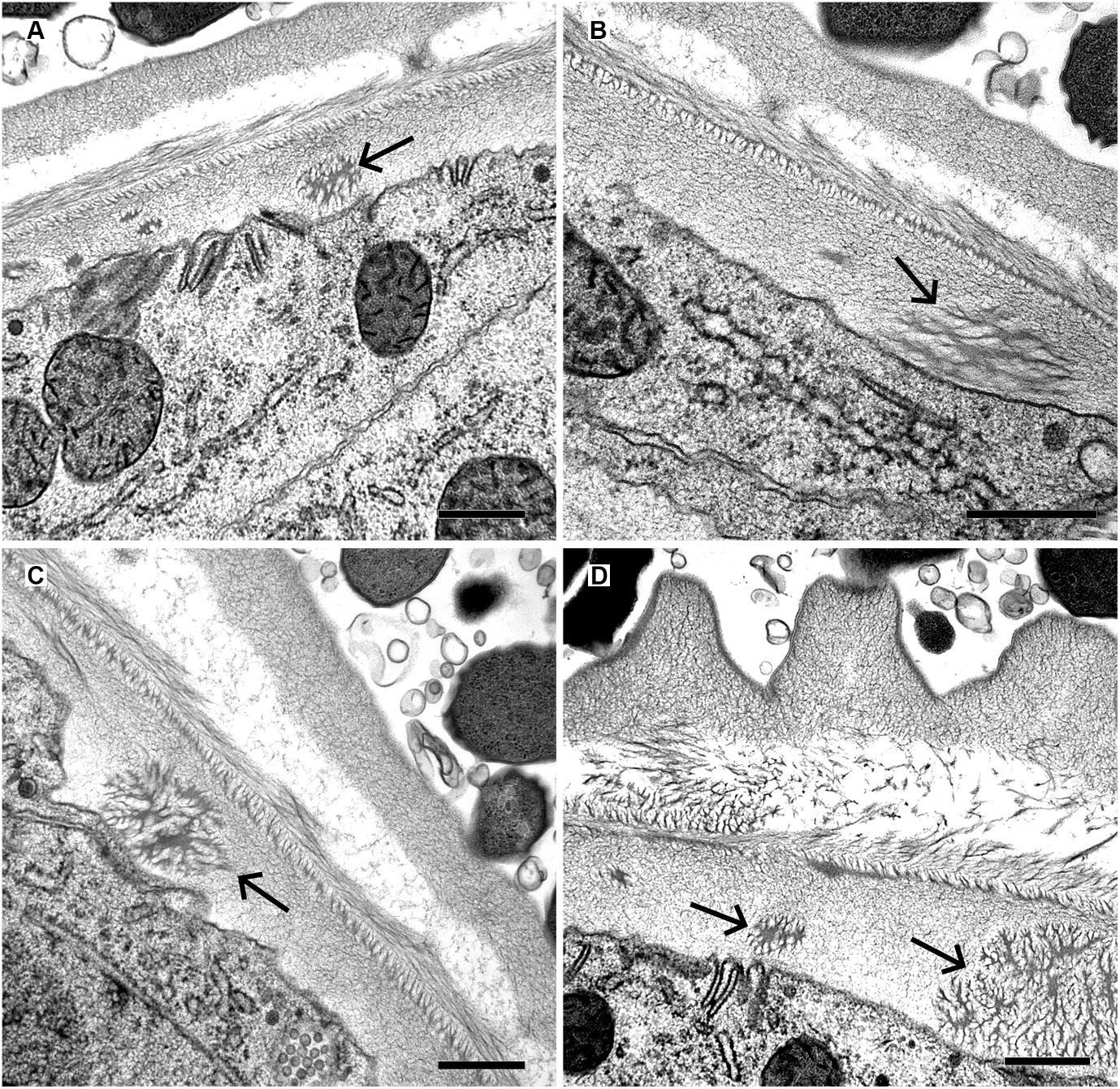

**Figure EV5 - The cuticle structure of *gcp-2.2* knockout strain of *C. elegans*.**

Several views on unspecific spongy structures situated in the basal layer of the cuticle (arrows). Scale bar shows 500 nm.
