## Appendix for "Uncovering the essential roles of human GCP 2 orthologs in *Caenorhabditis elegans*"

* equal author contribution

Table of content	page

Appendix Figure S1…………………………………………………………………………....2

Appendix Figure S2…………………………………………………………………………....3

Appendix Figure S3…………………………………………………………………………....4

Appendix Figure S4………………………………………………………………………....5, 6

Appendix Table S1…………………………………………………………………….……....7

Appendix Table S2…………………………………………………………………….……....8

Appendix Table S3…………………………………………………………………….……....9

Appendix Table S4…………………………………………………………………….……..10

Appendix Table S5…………………………………………….……………………………..11

Appendix Supplementary References …………………...…….……………………………..12

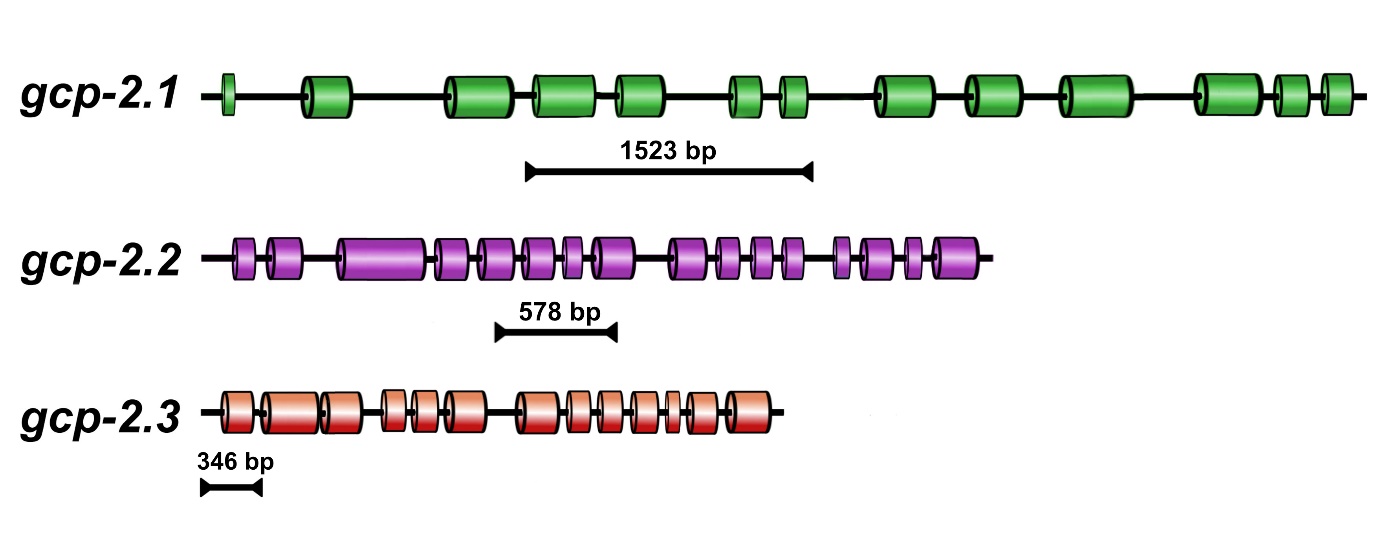

**Appendix Figure S1 – Scheme showing the deletion used for particular mutant worms.**

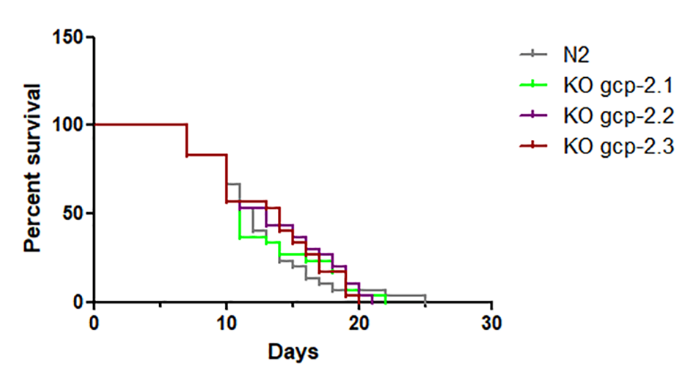

**Appendix Figure S2 - The effect of the gene knockout on the lifespan of *C. elegans* worms.**

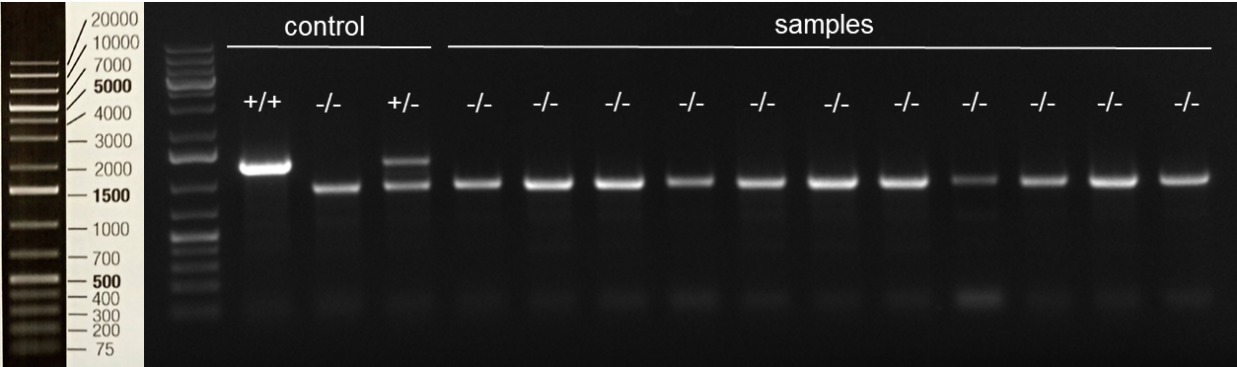

**Appendix Figure S3 – Gel after the backcrossing (*gcp-2.3* KO example).**

The indication in the figure is +/+ for wildtype, +/- for heterozygotes, and -/- for homozygous mutants.

***Caenorhabditis elegans, gcp-2.1* promoter area (2898 bp)**

ttttcaggaggacccttggaggctcaaccacttcatttatttatcttttttaagttcacc

cttcagcataccaaaggttacatttttgcagcagtgcacttataatgttgataataataa

aatattaattggaatatgatgtttgttatcttatcaggcaatagtcgacattgataaaaa

ttggaagattgctttcaaattgatttttttttccaattattttactttttgaaacatttc

attttacaattctattgttccttgagaaaaaaacgtcataagttttgattttttgcaatg

cgttcagttattttctgaacgaaacgtatttcatttgaaaggacttcttctcaatagtcc

actacaaataatttttatgcatattcaggtcaaatgtttattttatgaagtttcatagac

atggagacatttttatatctttcaagacagcctttcaaaaaattagtgaaattgtttaat

caagtccattaaaaatgttaaataagtaaaatgtttgtcgaattttttgaatgttttttt

tgagttactggagaatgaatcaaatgatttaaagtggtcgttttttacagtaaaaaagta

ttatattttgaatgactaatgtatttatttgataatataatttctttcagtaagatactt

tatgtgtcctatgcctttccacttctaaagccagaagtgaactgattggaaatttttttt

taagtttcaattttgtacccaaccgtagagttcgtatatacgagcattctcattaatttg

ccagtgtttcaaaacaaaaattccaagaaaaaaaaactctataattgcctcatttcttct

gttagtctggctgcttgaattttgaattataaatgttgactcaggacttattgccagccc

taattaaaaagagtattttttctattagcatggtatttgtattgcacacattttttacac

aattcctatgagattgaagttattaagcctttttgttacaattaaaataaaattgtcgaa

cttccatataaagtttaagtcttttcaagtaaaaaaatagtaagtccaaacacagaaaaa

tgttctgaaaattccctaataacggaatgacctttgaaaattagtattgtagccatacta

acaaaagaaatacggcaataatatcattataaaatattgtagtatagctataacgcaaat

accaaaccttcatcagtgtgtgtaatagaaagaacaagtgtcctactcaaacataccaat

aaacatacaacaataaggtgggcccagggagcgcacgtatcatatcctctttttctacac

atacaccagatgatggagcatcatcagacaactgacttcaattcgatgacactctggatt

atatgtatcaagagggggaaacaattgaggggagggggcagatgaaagaatcgtcgcttg

tgattctcaattcaattttcatttttctgtgtctgtttacaatattagtagagaaaagca

tttgatatatgaactaatgttttggggtttgtatcatagttttttaatcacaaaactgaa

aactttatttttatatattgttcatgcaaaaatgttagcatttccggttcatgatgtttt

ctatcaatttttgttgtgattacatttgcaaaattatttatttaatcttaaaggtggagt

agtgccagtgggaattttgtctaaatacacttattttgatcaaaaacgatcgaataccat

aatataacgttccaaaaattttttttagtgttttcataatttctgggcgaagttttggca

aattgccataattttgaaaaataagagcttttgaggaaatccaaagaaattacgcttatt

ccgagccctatgtacaatgtttcaatacaaataattaaaacaaaattgcactataaaaaa

taaaggaaaattttttttttggtcgactttcaaaatgagtggcaaaaacgaagtaattgt

cactttttgacagttgataaaaaatgttcaaaaaccttttgaaaagttttattgtgatct

ttggtcattttgggaccaaatgagtggtttataacaatttccccacttgcgctactccac

ctttaaaacaccaaaaagtgttaggctgttctatataatttgtgcccaaaaaatatgaca

tcagcatgttcttaacaatgaaaaatctgttgagaactctgagtctcttctcctgcattt

tttcaatagatctacgtagatcaaaccgaaatttttttcgtgcatttttttcagtcttag

aagggatagtttacttggcctaagaatacaaaaaatccacaattttgtttctataatcga

agaagtattgttttaaaatgcatgtatgtgatacaaatgaattaaattttaaacaaaatt

gaaaaattactgttttgaaacctgcaaatcttatctggcctaaatcaatgttgacagtcc

gtttacaatgagtttcatttttgttgctcagatatcaaatacacaattacacgtaaactt

ttgaaaagtgtattcatagtttcaaagttttcgttttttcttcgtccagtcatttcttac

agtaatttttgattaagttcttaactgtaactgtaattaattcaactccacttttaaata

atctcgcaacatttgtttgaaatgttgttgttccagaacaatagtttatatgtaaatgcc

gtacgtcggagttggagcacaaacagtttccacaagcttaacaggagcgcctatggtgaa

ggcatacattgcaattgctgcctctttgatttttgttttttgtattgctgcgttgggtgt

tcatcactcggaaagaaagttcaacaaattcaataaagtttcaattgatgacattcataa

atctgatgcagga

***Caenorhabditis elegans, gcp-2.2* promoter area (1107 bp)**

ttttcaggaggacccttggaggagagataattttcatttctatgattgtccgtttttgtt

cgttatgcataatgttttgactcctttcattacagtttgacacatttgtaaacgtttata

atcttcaataaacaacataagattttttttttgtaaaacctaacttataatctgacattt

tacgcttaaatcaggtttttttatttacggtaaatagaaagttttttcaacaaactaaac

ttgaagggggttttttttgcatcataagaaatggttctatacttcatcttggactttgat

atgaaatagtaatttttcgattaaagagtgttactgttagtaactgcaaccgctttcgtt

gcagcagagaaggttcagccatgaatgatgttttgaaatatcaaaactaaattcaatggg

aatccatgatctcaccaaataataacactctgaaccttccagtttgattatttttcagta

ttctataaaatactttttttatttcatttcaaacattcagtcgaataattgagcctgatt

ttcaaatgatccattatccccgagtttaagcagttttgagttgttaaagtcttgattatt

aatcaataaaaacaaagttgtattcgatctttgtttaatgtatctttcgtgctcagattt

tattaggacatatcatatatgagtttttatatttgaaaaaaaccaatcagtcacacacaa

ttttccatcgtcgtttatacttttatgggcgttgaaacgacaacccacattttcccaaaa

aattttcttccttgactactctgaatatatttctattaaattcatttttgtaaaatttga

tcctaatttaagtgtagaaaatgataacagctcggaaccggtttacacaaggttgtagtt

gtatgaagtattataacaataatggagacaaagtattccaaaagtctaaaagaaatagag

gaatgaaagctagcggtgtggttcttgttgcagtttctactgttgctttgactattattt

tatctaatgcgatacatcaatcttacaaatcaaattcaaagcctcttcctaaattatcaa

ttgtaccggtagaaaaaatgagtaaag

***Caenorhabditis elegans, gcp-2.3* promoter area (911 bp)**

ttttcaggaggacccttggaggtcacgtttgatagactctcagagtaataattatttaaa

tgtcaactttttaaagtgtaaggtggagtagtgcaagtggggaaagtgtttaaatggtga

aaatgaccaaaaataattgcaaaacattacaacaaaattttggaatgtttttatttactg

caaaaaaatgatacctactcagtttttgccagtgcgacataagtctggatttgcttgaaa

gcttacagttacagaaattttttaaaaactcttgaattttggagtgttttattgcgatat

tcagacaaaatccccactggcgatgtttactatttaaccaataaatgtattttgaataca

gtaacgttgagaaaacttgacattaaaaaaactcacagaaaatttcgaattagtttttca

tttgacccgttcgaatgtacattcgttcaaatttcaaccaacacattccaacggaagttt

gatgcctgttcaaaaatcatggacagattatttttttaatcaggaacgagagcatgtcga

acacatctaacgccgaaaactagtttttatcaggaacttctgaattaaatccctaaaata

atgtattcttacgaataaataatgaaactattttttaaactcgtcgactttgccagtaat

atttgcattttatcaaagcatataaatataacctggtaaacgaatcacttttctatgata

taacgtttttcaaaaatcgaacatgaaaaaaggattacaaatattcggcgttgtattact

tctagcagcgacagttgttgtaactgtactcatttcaaattatgtacatcagctttcaat

gtcgagtgggacacctacaatacaaaatacagtttcaatcgcaaatgtaccggtagaaaa

aatgagtaaag

**Appendix Figure S4 – Promoter region for *gcp-2.1*, *gcp-2.2* and *gcp-2.3*.**

**Appendix Table S1 - Statistical evaluation of the data for the impact of gene knockouts on the reproduction of the worm (Fig. 7 A in the article)**

| Kruskal-Wallis test | |
| --- | --- |
| P value | 0.0054 |
| Exact or approximate P value? | Gaussian Approximation |
| P value summary | ** |
| Do the medians vary signif. (P < 0.05) | Yes |
| Number of groups | 4 |
| Kruskal-Wallis statistic | 12.69 |

| Dunn's Multiple Comparison Test Summary | Difference in rank sum | Significant? P < 0.05? | Summary |
| --- | --- | --- | --- |
| N2 vs KO gcp-2.1 | 7.600 | No | ns |
| N2 vs KO gcp-2.2 | 12.80 | Yes | ** |
| N2 vs KO gcp-2.3 | 9.600 | No | ns |
| KO gcp-2.1 vs KO gcp-2.2 | 5.200 | No | ns |
| KO gcp-2.1 vs KO gcp-2.3 | 2.000 | No | ns |
| KO gcp-2.2 vs KO gcp-2.3 | -3.200 | No | ns |

Explanation of characters used in the tables. P value < 0.001 - Extremely significant (***); P value 0.001 to 0.01 - Very significant (**); P value 0.01 to 0.05 – Significant (*);

P value > 0.05 - Not significant (ns).

**Table S2 - Statistical evaluation of the data for the impact of gene knockouts on the pharyngeal pumping (Fig. 7 B in the article)**

| Kruskal-Wallis test | |
| --- | --- |
| P value | < 0.0001 |
| Exact or approximate P value? | Gaussian Approximation |
| P value summary | **** |
| Do the medians vary signif. (P < 0.05) | Yes |
| Number of groups | 4 |
| Kruskal-Wallis statistic | 30.12 |

| Dunn's Multiple Comparison Test Summary | Difference in rank sum | Significant? P < 0.05? | Summary |
| --- | --- | --- | --- |
| N2 vs KO gcp-2.1 | -22.60 | Yes | *** |
| N2 vs KO gcp-2.2 | -12.15 | No | ns |
| N2 vs KO gcp-2.3 | 2.750 | No | ns |
| KO gcp-2.1 vs KO gcp-2.2 | 10.45 | No | ns |
| KO gcp-2.1 vs KO gcp-2.3 | 25.35 | Yes | *** |
| KO gcp-2.2 vs KO gcp-2.3 | 14.90 | Yes | * |

Explanation of characters used in the tables. P value < 0.001 - Extremely significant (***); P value 0.001 to 0.01 - Very significant (**); P value 0.01 to 0.05 – Significant (*);

P value > 0.05 - Not significant (ns).

**Appendix Table S3 - *C. elegans* strains used in the study.**

| Gene | Strain name | Genotype/Allele Name  (WormBase) | Sequence Name | Source |
| --- | --- | --- | --- | --- |
|  | N2 | *C. elegans* wild isolate |  | CGC^1^ |
| *gcp-2.1* | RB1055 | ok1004 | R57.1 | CGC^1^ |
| *gcp-2.2* | Gcp-2.2 | tm6541 | C35C5.2 | Dr. Mitani/NBRP^2^ |
| *gcp-2.3* | C35C5.11 | tm5414 | C35C5.11 | Dr. Mitani/NBRP^2^ |

^1^ *Caenorhabditis* Genetics Center, University of Minnesota, MN, (Ann E. Rougvie)

^2^ National Bioresource Project for the Experimental Animal “Nematode *C. elegans*”

**Appendix Table S4 – Primers designed for genotyping after backcrossing and for amplification of promoter regions of *gcp-2.1*, *gcp-2.2* and *gcp-2.3*** (restriction sites underlined).

| **Primers for genotyping after backcrossing** | |
| --- | --- |
| ***gcp-2.1*** | |
| R57_lex3-F | AAATGCGTGGTCGGAAGCAG |
| R57_1ex5-R | TCTGAACACCCTCATTAGGC |
| R57_1ex8-R | GATCCATAGACCCATGCGTC |
| ***gcp-2.2*** | |
| genGCP2.2IntFwd | ACGTTGACAACATTCGCTCC |
| genGCP2.2IntRev | ACATCGCCTGAGTTTCTAGT |
| ***gcp-2.3*** | |
| genGCP2.3ExtFwd | CAACGGAAGTTTGATGCCTG |
| genGCP2.3ExtRev | GAAGGGAATGCCGGCAATCT |
| **Primers for amplification of promotor regions** | |
| ***gcp-2.1*** | |
| R57proF | AAAAAAGCTTCTCAACCACTTCATTTATTTATC |
| R57proR | AAAAGGATCCTTACATATAAACTATTGTTCTGGAAC |
|  | restriction sites – HindIII and BamHI |
| ***gcp-2.2*** |  |
| FRD2.2promBamHI | GATAGGATCCAGAGATAATTTTCATTTCTATGATTGTCC |
| REV2.2promKpnI | GTAAGGTACCTTTCTACACTTAAATTAGGATCAAATTTTAC |
|  | restriction sites – BamHI and KpnI |
| ***gcp-2.3*** |  |
| FRD2.3promBamHI | GATAGGATCCTCACGTTTGATAGACTCTCAGAGTAA |
| REV2.3promKpnI | GTAAGGTACCGTTCGATTTTTGAAAAACGTTATATC |
|  | restriction sites – BamHI and KpnI |

**Appendix Table S5 – PCR protocol for analyses of homozygous or heterozygous lines of KO worms after backcrossing**.

**KO *gcp-2.1***

**PCR mix:**

| Ready To Use PCR MasterMix 12 | 12.5 µl |
| --- | --- |
| R57_1ex3-F | 0.5 µl |
| R57_1ex5-R | 0.5 µl |
| R57_1ex8-R | 0.5 µl |
| DNA template | 1 µl |
| PCR-grade water | 10 µl |

**cycler setting:**

| initial denaturation | 3 min | 94 °C | 1x |
| --- | --- | --- | --- |
| denaturation | 30 sec | 94 °C | 40x |
| annealing | 1 min | 53 °C | 40x |
| extension | 3 min | 72 °C | 40x |
| final extension | 5 min | 72 °C | 1x |

**KO *gcp-2.2***

**PCR mix:**

| Ready To Use PCR MasterMix 12 | 12.5 µl |
| --- | --- |
| genGCP2.2IntFwd | 0.5 µl |
| genGCP2.2IntRev | 0.5 µl |
| DNA template | 1 µl |
| PCR-grade water | 10.5 µl |

**cycler setting:**

| initial denaturation | 3 min | 94 °C | 1x |
| --- | --- | --- | --- |
| denaturation | 30 sec | 94 °C | 35x |
| annealing | 1 min | 53 °C | 35x |
| extension | 2 min | 72 °C | 35x |
| final extension | 5 min | 72 °C | 1x |

**KO *gcp-2.3***

**PCR mix:**

| Ready To Use PCR MasterMix 12 | 12.5 µl |
| --- | --- |
| genGCP2.3ExtFwd | 0.5 µl |
| genGCP2.3ExtRev | 0.5 µl |
| DNA template | 0.5 µl |
| PCR-grade water | 11 µl |

**cycler setting:**

| initial denaturation | 3 min | 94 °C | 1x |
| --- | --- | --- | --- |
| denaturation | 30 sec | 94 °C | 35x |
| annealing | 1 min | 58.5 °C | 35x |
| extension | 2 min | 72 °C | 35x |
| final extension | 5 min | 72 °C | 1x |

**Appendix Supplementary References – References to „Figure EV1 –** **Primary sequence alignment of three *C. elegans* GCP2.1 splice variants“.**

- Davis P, Zarowiecki M, Arnaboldi V, Becerra A, Cain S, Chan J, Chen WJ, Cho J, da Veiga Beltrame E, Diamantakis S, *et al* (2022) WormBase in 2022—data, processes, and tools for analyzing Caenorhabditis elegans. *Genetics* 220
- Sievers F, Wilm A, Dineen D, Gibson TJ, Karplus K, Li W, Lopez R, McWilliam H, Remmert M, Söding J, *et al* (2011) Fast, scalable generation of high‐quality protein multiple sequence alignments using Clustal Omega. *Molecular Systems Biology* 7: 539
